# Closed-loop brain stimulation to reduce pathologic fear

**DOI:** 10.1101/2022.07.24.501314

**Authors:** Rodrigo Ordoñez Sierra, Lizeth Katherine Pedraza, Lívia Barcsai, Andrea Pejin, Gábor Kozák, Yuichi Takeuchi, Magor L. Lőrincz, Orrin Devinsky, György Buzsáki, Antal Berényi

**Affiliations:** MTA-SZTE ‘Momentum’ Oscillatory Neuronal Networks Research Group, Department of Physiology, University of Szeged, Szeged, Hungary; Department of Biopharmaceutical Sciences and Pharmacy, Faculty of Pharmaceutical Sciences, Hokkaido University, Sapporo, Japan; Department of Physiology, Anatomy and Neuroscience, Faculty of Sciences University of Szeged; Szeged, 6726, Hungary; Neuroscience Division, Cardiff University, Museum Avenue, Cardiff CF10 3AX, UK; Department of Neurology, NYU Langone Comprehensive Epilepsy Center, NYU Grossman School of Medicine, New York, NY 10016, USA; Neuroscience Institute, New York University; New York, NY 10016, USA; Center for Neural Science, New York University, New York, NY 10016, USA; HCEMM-SZTE Magnetotherapeutics Research Group, University of Szeged; Szeged, 6720, Hungary

## Abstract

Maladaptive processing of trauma related memory engrams leads to dysregulated fear reactions. In post-traumatic stress disorder (PTSD), dysfunctional extinction learning prevents discretization of trauma-related memory engrams and leads to generalized fear responses. PTSD is postulated as a mnemonic-based disorder, but we lack markers or treatments targeting pathological fear memory processing. Hippocampal sharp wave-ripples (SWRs) and concurrent neocortical oscillations are scaffolds to consolidate contextual memory, but their role during fear processing remains poorly understood. We demonstrate that closed-loop SWRs triggered neuromodulation of the medial forebrain bundle (MFB) can enhance the consolidation of fear extinction. It modified fear memories that became resistant to induced recall (i.e., ‘renewal’ and ‘reinstatement’) and did not reemerge spontaneously as a PTSD-like phenotype. The effects are mediated by D2 receptor signaling induced synaptic remodeling in the basolateral amygdala. These results suggest that SWRs help consolidating fear extinction memories. Furthermore, enhancing the consolidation of extinction engrams by SWR-triggered induction of reward signals can alleviate pathologic fear reactions in a rodent model of PSTD. No adverse effects were seen, suggesting this potential therapy for PTSD and anxiety disorders.

## INTRODUCTION

Posttraumatic stress disorder (PTSD) is a debilitating psychiatric disorder resulting from direct or indirect stressors, threats or life-threatening events perceived to compromise personal physical or mental safety^1-3^. Symptoms include intense feelings of unprovoked fear, panic attacks, anxiety, intrusive fear memories during wakefulness or in nightmares, fear generalization and avoiding similar but neutral stimuli^4, 5^. PTSD is highly resistant to psycho- and pharmacotherapy ^6-8^.

Experimental and clinical studies revealed altered memory formation resistant to normal processes of extinction as core PTSD features^9-12^. Memory alterations include involuntary hypermnesia or explicit amnesia for trauma-related stimuli and fear generalization to non-trauma related stimuli in animal models and human patients^13-15^. Novel models explore how pathological fear memories are consolidated^16-18^, extinguished^19-22^ and reconsolidated^23-26^.

Learning unpleasant things and remember them is advantageous for the organism for avoiding future reoccurrences. Irrelevant memories fade away either by graceful degradation^27, 28^ or by another type of learning called active extinction^29, 30^. Paradoxically, these two types of memory consolidation processes compete with each other, perhaps with different mechanisms, and different behavioral consequences. Current models conceive PTSD as mnemonic-based, but we lack the mechanistic understanding of pathological memory consolidation^31^. Impaired extinction may fail to extinguish traumatic memory leading to their intrusion in inappropriate contexts and, thus, become maladaptive.

Hippocampal sharp wave ripples (SWRs) are a rich source of systemic and local information underlying memory consolidation in normal and pathological conditions^32^. Disrupting SWRs can impair performance^33-35^. Long-duration ripples predominate after successful acquisition of memory in a hippocampus-dependent task and optogenetic prolongation of spontaneous ripples enhances memory consolidation^36^. SWRs promote the structured ‘replay’ of hippocampal place cells’ activity patterns following learning^37-40^. SWR-triggered activation of the internal reward systems during hippocampal replay can effectively induce new explicit memory traces^41^. A fraction of CA1 place cells are engram neurons of contextual contingencies beyond spatial localization^42^, and CA2 pyramidal neurons active during a social recognition task can be reactivated during SWRs^43^. The molecular, cellular and oscillatory activity underlying hippocampal-dependent consolidation of fear memories are understood^44,46-51^, but the role of hippocampal SWRs during fear processing remains poorly understood.

Fear conditioning is a validated PTSD model in humans and animals^45-47^ and fear reduction achieved by exposure-based extinction procedures are context-dependent, suggesting that hippocampal representation of the extinction context drives fear attenuation^48^. Basolateral amygdala activity decreases with conditioning stimuli (CS+) when animals are exposed to the same context used for extinction, but increases following CS+ non-extinction exposure^49^. Hippocampal inactivation enhances extinction to CS+ promoting low fear expression in environments different from the extinction context^50, 51^. We hypothesize that facilitating the extinction of memories through manipulating internal reward signals during extinction learning may attenuate traumatic memories in inappropriate contexts, thus reducing pathologic fear reactions.

We found that SWRs help mediating fear extinction and that SWR-triggered closed-loop stimulation of the reward system medial-forebrain bundle (MFB) can enhance extinction of fearful memories. This reduced fear expression across different contexts and prevented excessive and persistent fear responses. The effect is mediated by BLA G protein Rac1 and D2 receptors. Selective suppression of SWRs after extinction delayed fear attenuation, suggesting that extinction learning requires intact SWRs. These findings highlight the prominent role of SWRs in fear extinction and suggest that closed-loop neuromodulation may reduce PTSD symptoms by targeting oscillatory activity related to memory processing.

## RESULTS

### SWR-driven closed-loop electrical stimulation of the medial forebrain bundle accelerates extinction and prevents fear recovery

Rats underwent a single session of fear conditioning to develop PTSD-like phenotypes (Supplemental Figure 1A-E). Fear extinction (i.e. twenty re-exposures per day in four blocks to CS+ in a new context without US) was performed on consecutive days until a remission criterion (reduction of freezing behavior to < 20 % of the initial freezing) was reached or up to maximum seven days (Fig. 1A). In one cohort of the animals, MFB was stimulated during SWRs in a closed loop manner (fourteen 1-ms long, 100μA square-wave pulses at 140 Hz) to assign a reward signal to the replayed extinction engrams. In another cohort of animals, stimulation was jittered in time (i.e. open loop animals). The third group received no electrical stimulation (control animals) (Fig. 1B). Fear related behavioral performance after extinction was tested by exposure to CS+ in hybrid context mixing new features with the conditioning context (‘RENEWAL TEST’) and by unpredictable exposure to the US (‘REINSTATEMENT TEST’). The persistence of the extinction was assessed by exposing the animals to CS+ 25 days following extinction (REMOTE TEST’).

**Figure 1.**
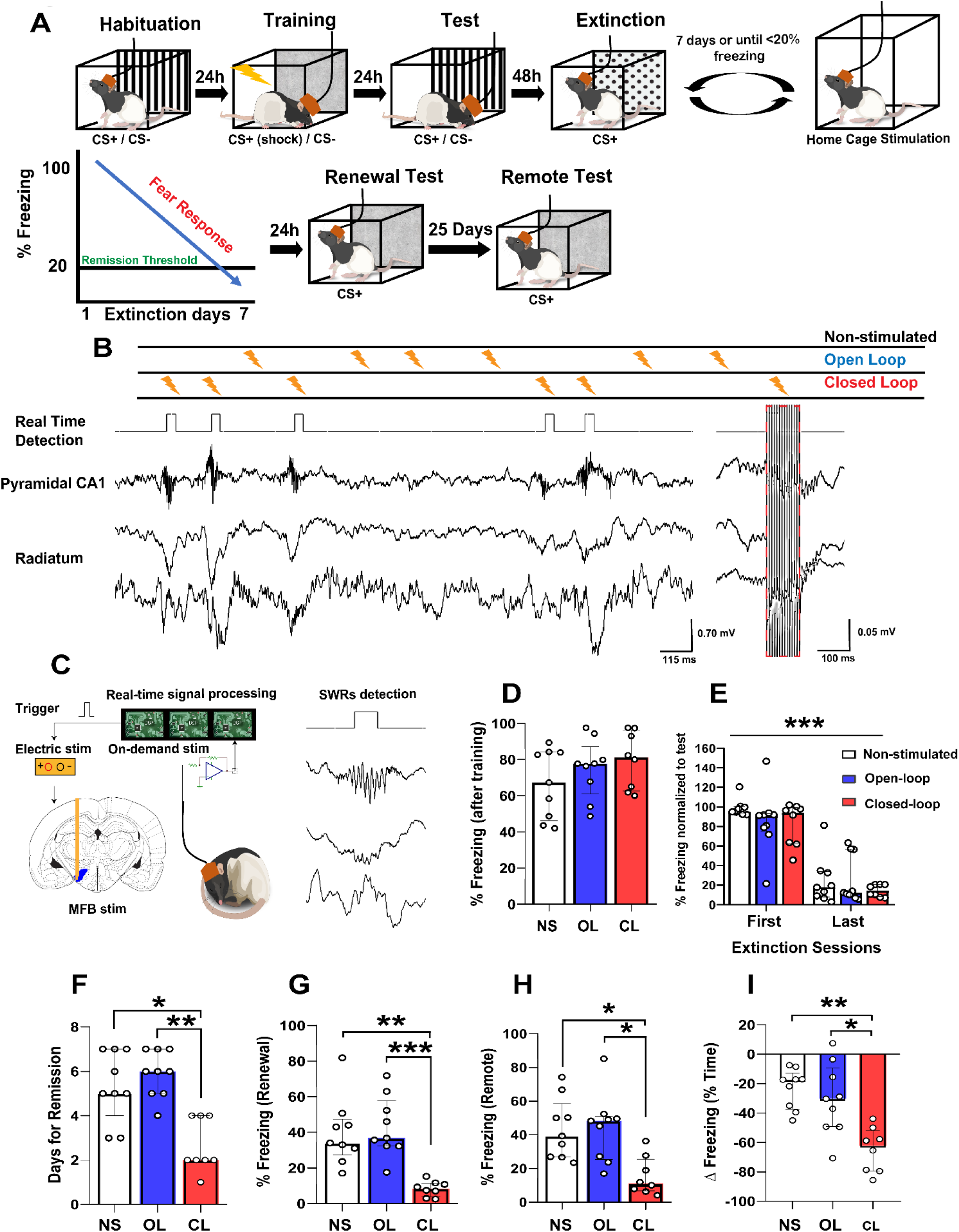
Closed-loop SWR-timed medial forebrain bundle electrical stimulation attenuates PTSD-like memories. **(A)** Schematics of the experimental design. Animals underwent fear conditioning training followed by a test session to evaluate memory recall, extinction sessions and 1h closed-loop stimulation where the online detected SWRs triggered MFB stimulation until achieving the remission criterion (reduction of freezing behavior to < 20 % of the initial freezing). One and twenty-five days after the last extinction session, renewal and fear recovery were assessed, respectively. **(B)** Closed-loop stimulation consisted of MFB stimulation during detected SWR events, open-loop stimulation was similar to closed-loop stimulation except jittered from SWRs (top). Representative LFP signals from dorsal hippocampus showing SWR events (the red lines represent detected SWRs) and stimulation pattern (red dashed rectangle; bottom). **(C)** A custom threshold crossing algorithm was used to trigger the MFB stimulation (fourteen 1-ms long, 100 μA square-wave pulses at 140 Hz) following SWR online detection. (**D)** No difference in fear expression in response to the CS+ following training between the three experimental groups (non-stimulated (NS) n=9; open-loop (OL) n=9; closed-loop (CL) n=8). € No difference between the fear expression of the three groups during the first 5 CS+ block after first and last extinction days. There was a significant decrease in fear expression over time, suggesting that extinction can attenuate fear. Values are normalized to the freezing expressed immediately after foot shock training (i.e. “Test”). (**F**) Animals exposed to closed-loop stimulation required less extinction sessions to achieve the remission criterion compared to the open-loop and non-stimulated groups. (**G**) Closed-loop neuromodulation induced lower fear expression during the renewal test in a hybrid context. (**H**) Closed-loop neuromodulation prevented spontaneous fear recovery 25 days after extinction. (**I)** Closed-loop neuromodulation reduces and maintains low fear expression 25 days following extinction. * = p< 0.05, ** = p< 0.01, *** = p< 0.001.

The rewarding properties of the MFB stimulation were verified using a conditioned place preference task (Supplemental Figure 1F). No significant differences were found in the fear expression between groups in the test after conditioning to CS+ (Fig. 1D) or after the first or the last extinction days (Fig. 1E). Supplemental Table 1 shows the results of descriptive and comparative statistics. Although extinction can overcome fear, animals exposed to closed-loop stimulation required less extinction sessions to achieve < 20 % of the initial freezing (i.e. the remission criterion) compared to the open-loop and non-stimulated groups (Fig. 1F). Following the exposure to the ‘renewal test’ in a hybrid context there was a significant decrease in fear expression in the closed-loop treated animals compared to the open-loop and non-stimulated groups (Fig. 1G). These results indicate that closed loop MFB stimulation during SWRs events can enhance fear extinction, decrease the time needed to achieve fear attenuation and maintain freezing levels low in challenging situations such as exposure to hybrid contexts resembling the learning contingencies.

To evaluate effect persistence, animals were exposed to a ‘remote test’ 25 days following the renewal in the hybrid context. Animals were kept in their home cages between the renewal and remote tests. Freezing in closed-loop stimulated animals remained at low levels compared to the open-loop and non-stimulated group (Fig. 1H), suggesting fear attenuation was resistant to spontaneous recovery.

Finally, we quantified Δ freezing as reduced fear reactions between those after fear condition and the remote test (Δ freezing = Freezing extinction – Freezing test CS+) to reveal the overall effect of the interventions (Supplemental Figure 1b shows the performance of individual animals in each group). Closed-loop simulated animals had stronger fear reduction than open-loop and non-stimulated animals (Fig. 1I). Together, closed-loop neuromodulation of the reward system triggered by memory consolidation related neuronal oscillations accelerates fear extinction and promotes persistent low fear expression of PTSD-like memories.

### Exploring non-specific effects and potential side effects of closed-loop MFB stimulation on memory functions

Since MFB stimulation is rewarding, we explored if stimulation without any extinction exposition reduced fear. After fear conditioning, animals received SWR-triggered closed-loop stimulation during sleep for three consecutive days but were not exposed to the extinction paradigm (Fig. 2A). The number of stimulation sessions were matched to the mean number of extinction sessions (i.e. 2.625 days) required in the previous experiment to achieve the remission criterion (Fig. 1F). Stimulation duration were identical to the previous experiment with extinction. The control group (non-stimulated; NS) was exposed to identical fear conditioning followed by spending three days in their home cage without any intervention. No significant differences were found between the two groups immediately after CS+ conditioning (Fig. 2B) nor after three days of stimulation sessions (Fig. 2C). Thus, the closed-loop SWR-triggered stimulation alone, without extinction, did not decrease fear expression.

**Fig 2.**
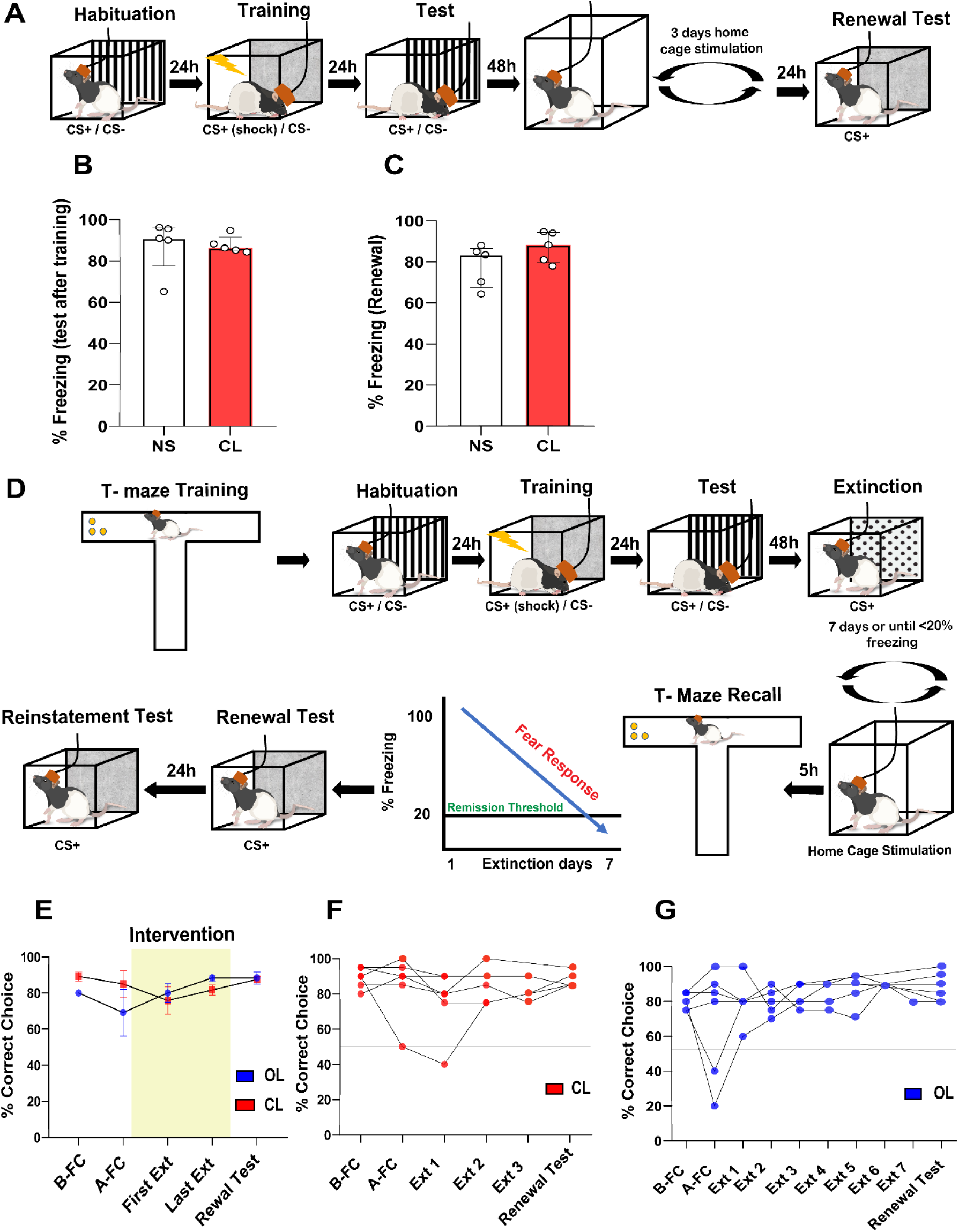
Contribution of fear extinction and side effects on co-storaged memories during closed-loop MFB stimulation. **(A)** Schematics of the experimental design. Fear conditioning and test was performed as before. Closed-loop animals were exposed to 3 consecutive SWR-triggered stimulation sessions without extinction. No difference was found in fear expression in response to the CS+ following training **(B)** and renewal **(C)** between the groups (non-stimulated (NS) n=5; closed loop (CL) n=5). **D)** Before fear conditioning, animals were trained in a visual cue forced alternation T-maze task until achieving 80% of correct choice. Next, animals were exposed to fear conditioning, extinction and stimulation following Fig 1 (open-loop (OL) n=6; closed-loop (CL) n=6). **(E)** T-maze performance was unaltered during the experiments regardless of the stimulation type. Individual performance of the animals is shown for open-loop **(F)** and closed-loop **(G)**.

We next tested if the SWR triggered closed-loop stimulation interferes with already consolidated non-fear related memories as a non-specific detrimental effect. Animals were trained in a spatial memory task, where a randomly alternated visual cue indicated the correct choice in a T-maze to receive reward (froot-loops pellet). A total of 20 trials per day were performed until achieving 80% of correct choice. Afterwards, fear conditioning, extinction and stimulation sessions were performed the same way as in the previous experiment until achieving remission (Fig 2D). They were also retested in the same spatial memory task each day during the extinction procedure. Extinction+stimulation sessions and T-maze were separated by five hours and the order of the behavioral tasks were randomized across the experiment. Both OL and CL stimulated animals maintained performance in the T-maze (Fig. 2E; individual performance during the fear conditioning and extinction procedure are showed in Fig. 2F-G).

These results suggest that 1) the beneficial effect of the closed-loop stimulation is not generic, but it enhances extinction learning, and 2) already consolidated non-traumatic memories are not affected by the stimulation.

### The enhancement of extinction induced by closed-loop stimulation is mediated by D2-receptor and G protein Rac1 in BLA

We next explored plasticity-dependent mechanisms induced by closed-loop MFB stimulation and resulting enhanced fear extinction. We tested the potential contributions of BLA dopamine receptors and the small G protein Rac1, a Rho family member involved in learning-induced synapse formation ^52-55^. Animals underwent the prior experimental protocol, but immediately after each extinction session and before the closed-loop stimulation, the BLA was bilaterally microinfused with the Rac1 inhibitor NSC2376, D1R antagonist SCH23390 or D2R antagonist sulpiride (Fig. 3A-B). No significant differences were found in the test after conditioning to CS+ (Fig. 3C), nor in the fear expression during the first 5 CS+ block from first and last extinction day (Fig. 3D). Animals co-infused with NSC2376 and sulpiride required more days to achieve extinction than controls, closed-loop stimulated animals and closed-loop stimulated animals infused with SCH23390 (Fig. 3E). During the renewal test in the hybrid context, only sulpiride suppressed closed-loop stimulation’s effect (Fig. 3F). Similar to the renewal test, animals infused with sulpiride showed a significant fear recovery after the exposition to an immediate foot-shock protocol (Fig. 3G).

**Figure 3.**
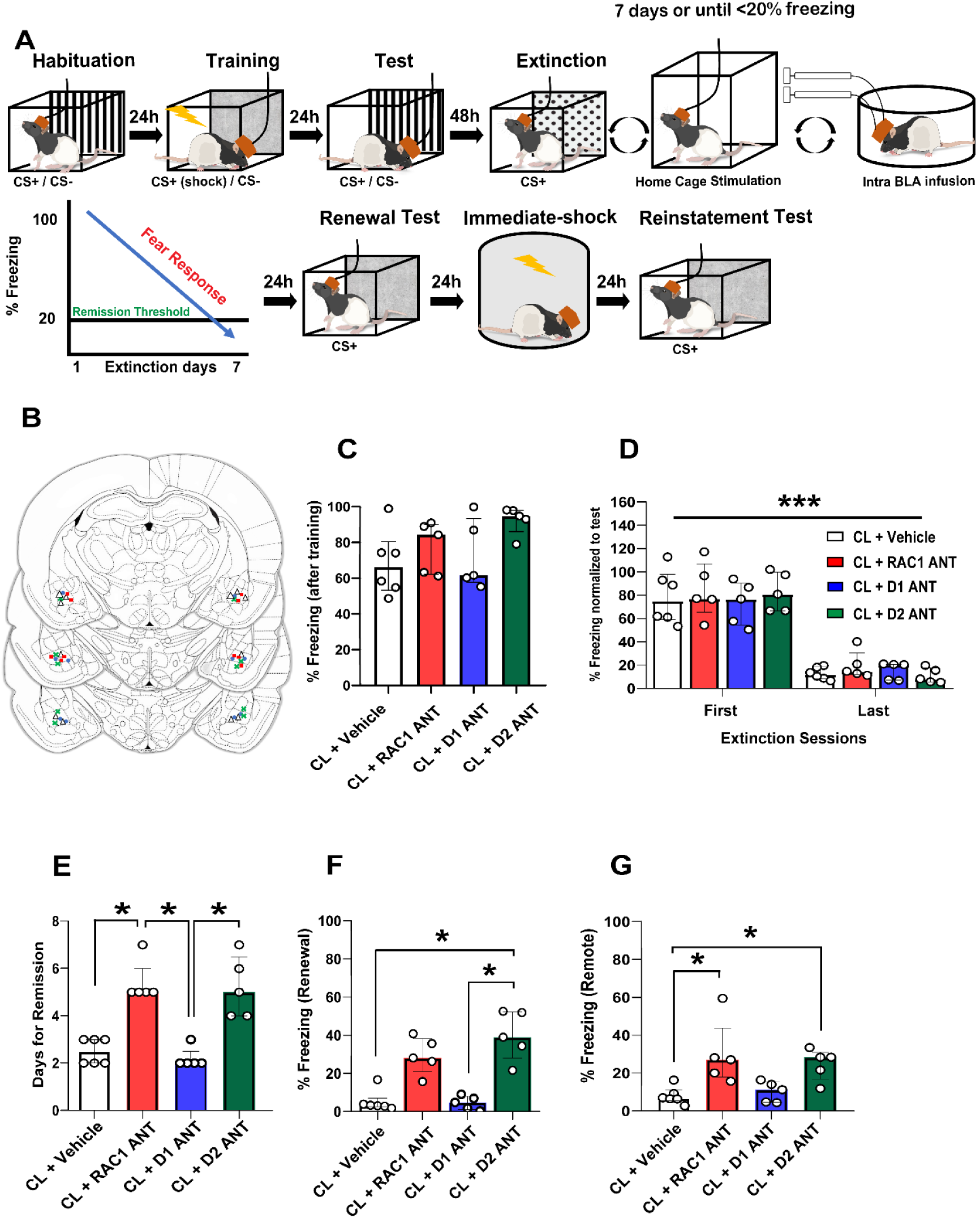
The closed-loop neuromodulation induced enhancement of extinction is mediated by Rac1 and D2Rs in the BLA. **(A)** Behavioral protocol and closed-loop neuromodulation was performed as before, in addition BLA was bilaterally microinfused with various drugs immediately after extinction and before home-cage stimulation. **(B)** Locations of cannula tips in each animal. Colors represent the different experimental groups. **(C)** No difference in fear expression in response to the CS+ following training between the four experimental groups (closed loop n=6; closed-loop+NSC2376 n=5; closed-loop+SCH23390 n=5; closed-loop+sulpiride n=5). **(D)** No difference between the fear expression of the four experimental groups during the first 5 CS+ block from first and last extinction day. Note the significant decrease in fear expression over time. (**E**) NSC2376 and sulpiride injected animals required more extinction sessions to achieve the extinction criterion. **(F)** Sulpiride suppress the extinction enhancement induced by closed-loop neuromodulation during renewal. (G) Animals treated with NSC2376 and sulpiride are more prone to fear recovery compared to the other groups. * = p< 0.05, *** = p< 0.001.

The pharmacological treatments did not modify the extinction criterion and fear processing without electrical stimulation (Supplemental Figure 2). Thus, NSC2376 and sulpiride prevented the enhancement of extinction induced by the closed-loop neuromodulation. This suggests that the closed loop neuromodulation-induced fear extinction involves dendritic spine plasticity mediated by RAC1 signaling and D2Rs in the BLA.

### SWRs are required to consolidate fear extinction

As fear extinction is a highly context-dependent process,^40-43^ and hippocampal SWRs are engaged in contextual memory consolidation across cortico-hippocampal circuits,^33, 56, 57^ we hypothesized that SWRs are required for fear extinction. To test this, we suppressed SWRs by ventral hippocampal commissural electrical stimulation that induces phasic silencing of hippocampal pyramidal cells and interneurons.^33, 34, 58^

Since some of the animals trained with high intensity foot-shocks resist extinction, we reduced the training intensity (US: 0.7 mA) to ensure that the extinction criterion was achieved within seven sessions in control conditions. During stimulation following each extinction, online detected SWRs triggered a single-pulse (0.5 ms) ventral hippocampal commissural stimulation (Fig. 4A-B). The stimulation intensity was adjusted for each animal to the minimal intensity required to disrupt the SWRs (range: 5–15 V). Open-loop animals were randomly stimulated within the same voltage range. No significant differences were found in the test after conditioning to CS+ (Fig. 4C) or in fear expression between groups during the first 5 CS+ block from first and last extinction day (Fig. 4D). SWR disrupted animals required more extinction sessions to achieve 80% of freezing reduction compared to open-loop animals (Fig. 4E) and expressed elevated levels of freezing in the hybrid context (renewal test) compared to the non-stimulated and open-loop groups (Fig. 4F). No differences were detected during the reinstatement test (Fig. 4G). These results suggest that hippocampal SWRs are required to consolidate fear extinction. The disruption of SWRs results in slow extinction learning and fear persistence in different environments beyond the extinction context.

**Figure 4.**
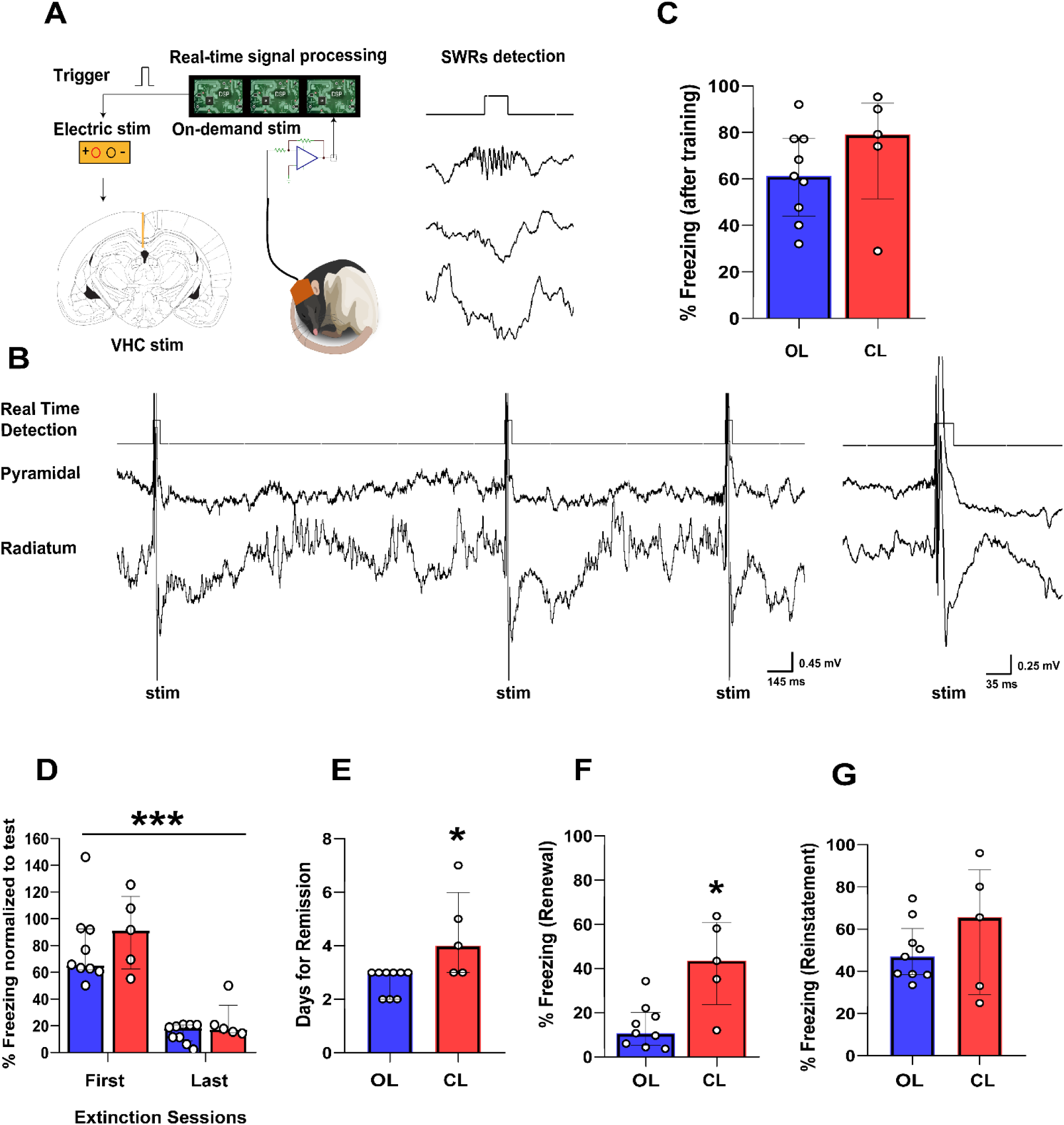
SWRs are required for the extinction of fear memories. **(A)** Behavioral protocol was performed as before, but SWR triggered VHC stimulation was performed for 1 h following each extinction session. **(B)** Representative LFP signals from dorsal hippocampus showing intact and disrupted SWRs events (red trace: detected SWRs, blue dots: timing of stimulation. **(C)** No difference in fear expression in response to the CS+ following training between the three experimental groups (open-loop (OL) n=9; closed-loop (CL) n=5). **(D)** No difference between the fear expression of the three experimental groups during the first 5 CS+ block from first and last extinction day. Note the significant decrease in fear expression over time. **(E)** SWR disrupted animals require more days to achieve the extinction criterion. **(F)** SWR disrupted animals show high fear expression during renewal. **(G)** No difference in fear expression during reinstatement. * = p< 0.05, *** = p< 0.001.

## DISCUSSION

Closed-loop stimulation of the MFB during SWRs enhance extinction of cued fear conditioning. SWR independent stimulation or stimulation without extinction learning was ineffective. Our intervention shortened the time to reduce fear expression. The effect persisted, since animals were resistant to both induced renewal, reinstatement and to the spontaneous reemergence of PTSD-like phenotypes even 25 days after treatment. These effects were mediated by D2 receptors and RAC1 signaling in the BLA, suggesting that closed-loop modulation of the reward pathways promotes a plasticity-dependent mechanism leading to extinction. Since disruption of SWRs increases extinction sessions required to achieve remission and predisposes animals to recurrent expression of pathological fear, the SWRs appear essential for extinction learning. These results offer novel avenues to develop closed-loop neuromodulation technologies for PTSD and anxiety disorders.

SWRs encode and consolidate spatial memory, and are involved in fear memory processing. Selective pre-or post-training inactivation of CA3 disrupts the acquisition and consolidation of contextual fear memory by reducing the number and dominant frequency of CA1 ripples and shifting underlying CA1 ensemble activity^59^. SWRs rely on synchronous CA1 principal neuron activation mainly controlled by PV+ interneurons^60^. Boosting the activity of hippocampal PV+ interneurons results in selective extinction of contextual fear memory and increased SWRs incidence^61^. Suppression of hippocampal PV+ interneurons results in altered principal neuronal phase coupling to SWRs, decreased ripple-spindle coupling and decreased consolidation of contextual fear memory^62, 63^. Our findings indicate that SWRs are required for the extinction of cued fear conditioning and can update the memory trace with rewarding information. Closed-loop disruption of SWRs delayed but did not block extinction since 80% of animals still achieved the remission criterion, consistent with the contextual dependence of fear extinction^49-51^, although cued fear conditioning is amygdala-dependent^64-66^. Our initial hypothesis that SWRs encode contextual features of ‘safety’ during the extinction is consistent with the decreased time to achieve remission but fails to explain the fear reduction to CS+.

SWRs are spatiotemporally precise windows to integrate information in neocortical and subcortical structures. A widespread increase in neocortical activity precedes SWRs^32^. We posit that during SWRs, replay and information integration involve contextual features of an engram as well as corresponding memory traces. This is supported by multiple roles of SWRs and hippocampal place cells in processing contingencies beyond spatial localization^42, 43^. Thus, the SWR triggered closed-loop MFB stimulation, and the resulting reward signal is coincident with a widespread ongoing brain network activity orchestrating the consolidation of fear extinction^67^ during SWRs events. Neuronal activity in the BLA increased during SWRs^68, 69^. The coordinated reactivation between the dorsal hippocampus and BLA during off-line aversive memory processing peaks around the SWRs^70^. The SWR-triggered closed-loop neuromodulation may provide a reward system safety signal to a consolidated aversive memory^71^ and/or enhances the network activity that encodes fear extinction^72^. In both cases, this potential mechanism resembles a counterconditioning process of memory updating using contrasting emotional valence^73-76^ with high temporal and neurochemical precision. This hypothesis is supported by the absence of closed-loop effect when animals are not exposed to the extinction learning. Under these circumstances the reward signal triggered by MFB stimulation is not coincident with SWRs activity promoted by extinction, preventing the enhancement of fear attenuation.

MFB fibers interconnect nodes critical for reward and emotional processing. The VTA sends dopaminergic axons to the NAc, amygdala and PFC via the MFB^77^. A cluster of dopaminergic neurons in the anterior VTA/SNc directly connect with CA1^78^. A global manipulation of the reward system through MFB deep brain stimulation may treat depression in animal models and human patients^79^. We found that temporally precise dopamine release in these circuits during SWRs may scaffold the extinction enhancement with BLA D2 receptors mediating the effects.

Multiple evidentiary lines support that dopamine released in the BLA during fear learning is controlling the saliency of the foot shock and the extinction through prediction error signaling of non-reinforced CS+ presentation^80^. Fear memories and extinction are encoded by different BLA neuronal populations. Thus, instead of overwriting, the extinction engrams can suppress the activity of neurons initially engaged in fear learning. Since neurons mediating extinction overlap with those responding to reward, activation of neurons that mediate extinction learning could also signal reward^81^. Our experimental design cannot differentiate whether post-extinction SWRs are related to the reactivation of the original fear memory or represent the consolidation of the extinction. However, increased dopamine release during SWRs could change the emotional valence of an engram replay or directly suppress neurons engaged in fear learning. Reward-responsive VTA neuronal activity is coupled to SWRs during quiet wakefulness^82^, supporting that dopamine release is modulated by SWRs. As dopaminergic projections from VTA innervate D2 expressing PV+ interneurons and suppresses principal BLA neurons, locally suppressing GABA release^83^. The suppression of feed-forward inhibition can induce LTP at excitatory afferent synapses in the BLA, an effect also mediated by D2 receptors^84^.

Dopamine stimulation of engram cells may enhance forgetting by activating Rac1/Cofilin, which modulates actin cytoskeleton and cellular morphology^28^. Inhibition of Rac1 activity in the dHPC impairs extinction of contextual fear memories^85^ and photoactivation of Rac1 in the motor cortex suppresses motor learning^54^.

Our findings suggest three sequential mechanisms underpinning closed-loop extinction enhancement: 1) SWRs reactivate the memory engram and memory trace in BLA. 2) MFB stimulation promotes dopamine release in BLA. 3) BLA dopamine release can induce D2 receptor mediated plasticity processes culminating in Rac1 activation. Blocking RAC1 signaling prevents spontaneous or closed-loop neuromodulation induced extinction. RAC1 inhibition without closed-loop neuromodulation did not prolong the number of sessions required for successful fear extinction. However, chronic treatment impairs expression of the extinction memory during renewal. Additional work is required to determine the mechanisms of interaction between dopamine receptors and RAC1 modulation.

Our results suggest a novel translational treatment of fear-related disorders. The US Food and Drug Administration (FDA) approved MFB stimulation for treatment-resistant depression in clinical trials, with promising efficacy^79, 86^. Although SWRs detection was invasive, a non-invasive method could use cortical slow-waves and spindles that concur with SWRs in animals^56, 87, 88^. Thus, closed-loop stimulation triggered by cortical EEG activity could replace SWRs detection. Further, non-invasive techniques (e.g., tDCS, TMS) could stimulate reward-associated cortical areas instead of penetrating electrodes.

Our new framework to study and treat fear-related disorders relies on closed-loop stimulation guided by classical biomarkers of memory consolidation. Temporally precise manipulation of the reward system during SWRs overcomes the resistance to extinction in an animal PTSD model. SWRs are critical for extinction learning. Although dopaminergic agonists can enhance fear extinction^20, 89^, our intervention avoids side effects with systemic treatments. (e.g., psychosis, pathological gambling). Our data suggest that relationship between SWRs, slow-waves and cortical spindles may offer a potential non-invasive therapy.

## MATERIALS AND METHODS

### Animals

Rats (120adult male Long-Evans, 300-450 g, 3-6 months old) were kept in a 12-hour light/ dark cycle. All experiments were performed in accordance with the European Union guidelines (2003/65/CE) and the National Institutes of Health Guidelines for the Care and Use of Animals for Experimental Procedures. The experimental protocols were approved by the Ethical Committee for Animal Research at the Albert Szent-Györgyi Medical and Pharmaceutical Center of the University of Szeged (XIV/218/2016).

### Surgery

The animals were anesthetized with 2% isoflurane and craniotomies performed according to stereotaxic coordinates. Intracortical electrode triplets (interwire spacing, 0.2-0.4 mm) (Kozák et al., 2018) targeting the anterior cingulate cortex (ACC) (AP: +1.0, ML: 0.5, DV: 1.4,), bilateral BLA (AP: -2.8, ML: 4.6, DV: 8.1 mm from the dura) and the bilateral CA1 subfield of the dorsal hippocampus (AP: -3.5, -4.5 and -5.5, ML: 2.0, 3.0 and 4.0, DV: 2.9 and 3.0 all mm from Bregma). To improve DH-SWRs detection, a custom-built microdrive (Vandecasteele et al., 2012) was used in some experiments, allowing the vertical adjustment over the CA1 subfield. A custom-built bipolar electrode consisting of two insulated (except 200 μm at the tip) Tungsten wires (interwire spacing, 0.4 mm) was implanted in the left medial forebrain bundle (AP: -2.8, ML: 2.0 mm, DV: 8.1 all mm from Bregma). LFP electrodes and the base of the microdrive were secured to the skull with dental acrylic (Unifast Trad, USA). Two stainless-steel screws above the cerebellum served as ground and reference for the recordings, respectively. A Faraday cage was built using copper mesh and dental acrylic on the skull around the implanted electrodes.

In experiments involving concomitant electrophysiological recording and local pharmacological infusion, in addition to electrodes, rats were bilaterally implanted with 25-gauge guide cannulas above the BLA (AP: -2.8, ML: 4.7, DV: 6.9 all mm from Bregma). Cannulae were fixed to the skull with dental acrylic (Unifast Trad). Caps were used to cover cannulae to avoid any accidental occlusion.

### Electrophysiological recordings and stimulation

Rats were housed individually in plastic home cages. LFP recordings were performed in the home cage and the fear conditioning box (see below). For home-cage recordings, walls of clear Plexiglas (42 × 38 cm, 18 cm tall) were incorporated allowing the normal functioning of the recording systems and animal movement. To avoid any twisting and over-tension of the recording cables, a bore-through electrical commutator (VSR-TC-15-12; Victory-Way Electronic) was used. Food and water were available *ad libitum*. All recording sessions took place in the same room using 12/12 h light/dark cycle with light onset/offset at 7h/19h The multiplexed signals were acquired at 500 Hz per channel for closed-loop neuromodulation experiments (Kozák and Berényi, 2017). The neuronal signals were preamplified(total gain 400X), multiplexed on head and stored after digitalization at 20-kHz sampling rate per channel (KJE1001, Amplipex, Szeged, Hungary). During home cage stimulation, preamplified signals were analyzed on-line by a programmable digital signal processor (RX-8, Tucker-Davis Technologies, Alachua, FL, USA) using a custom made sharp-wave ripple detection algorithm, as follows.

The LFP of pre-selected tripolar electrodes from CA1 pyramidal layer were demultiplexed and band-pass filtered (150–250 Hz), and RMS powers were calculated in real time for ripple detection. Threshold crossings triggered a stimulation train lasting 100 ms and composed of fourteen 1-ms long, 100μA square-wave pulses at 140 Hz) in the MFB or single pulse (5-15V in the ventral hippocampal commissure (VHC) (STG4008; Multi Channel Systems, Reutlingen, Germany) depending on the experiment performed. MFB stimulation was performed under current mode and VHC stimulation in voltage-controlled mode. The threshold of the detection algorithm was set for each rat separately. Behavioral (i.e. rewarding) effect of MFB stimulation was confirmed with a place preference task (see below).

### Drugs and infusions

The Rac1 inhibitor NSC2376 (10μg/μl), D1 dopamine receptor antagonist SCH23390 (0.50μg/μl) and D2 dopamine receptor antagonist sulpiride (1 μg/μl) were dissolved in sterile physiological saline (0.9% NaCl). NSC2376, SCH23390 and sulpiride were infused bilaterally into the BLA using a 33G gauge injectors connected to Hamilton syringes via 20-gauge plastic tubes. The infusion injectors tip protruding 2.0 mm below the tip of the cannula and aimed the BLA center. A total volume of 0.5 μl per side was infused by a microinfusion pump at a rate of 0.125 μl/min. Injectors were left in place for an additional minute to ensure proper drug diffusion. All drugs were infused after the extinction sessions.

### Auditory Fear Conditioning

The experiments were carried out in a fear conditioning apparatus comprising three contextual Plexiglas boxes (42 × 38 cm, 18 cm tall) placed within a soundproof chamber. Four different contextual configurations were used (Habituation and Test Context (A): square configuration, white walls with black vertical horizontal lines, white smooth floor, washed with 70% ethanol; Training Context (B): square configuration, grey walls, metal grid on black floor, washed 30% ethanol; Extinction Context (C): rectangular configuration, white walls with black dots, white smooth floor; and Renewal and Remote/Reinstatement context (D): hybrid context comprising a square configuration, grey walls from training context, white smooth floor, washed with 70% ethanol. All sessions were controlled using a MATLAB custom script.

#### Habituation

On day 1, animals were exposed to the habituation session in context A. After 2 min of contextual habituation, they were exposed to 5 alternating presentations of two different tones (2.5 or 7.5 kHz, 85 dB, 30 s). Tone time intervals were randomized (30-40 s) during the session. No behavioral differences were detected under exposition to the two frequencies.

#### Training

On day 2, cue fear conditioning was performed in context B. After 2 min of contextual habituation, animals received 5 trials of one tone (CS+: 7.5 kHz) immediately followed by a 2s long footshock as unconditioned stimulus (US: 1.0 mA, 0.7 mA or 0.5 mA, depending on the experiment performed). The other tone (CS−: 2.5 kHz) was presented 5 times intermittently but never followed by the US.

#### Test

On day 3, animals underwent fear retrieval in context A. After 2 min of contextual habituation, rats were exposed to presentations of the CS+ or CS-in two different sessions. Each session consisted of a block of five tones. The order of the CS+ and the CS− in each session was randomized. Sessions were repeated every 4-6 h.

#### Extinction

In context C, from day 5 until reaching the remission criterion (see below), rats received extinction training consisting of twenty CS+ presentations without the US (unreinforced tones). Tones were repeated with randomized intervals (30-40 s) during the session.

#### Fear Remission from Extinction

We used an extinction threshold criterion to assess the efficacy of fear reduction after extinction sessions similar to^90^. The block of the first five tones during each extinction session was assessed to determine fear reduction level of the given day. Considering individual differences under fear conditioning^90-92^ fear reduction during extinction was expressed as a fraction of the percentage of freezing expressed during the CS+ test (Day 3) (% Freezing Reduction = Freezing extinction x 100 / Freezing test CS+). Fear remission was considered achieved when animals expressed ≥80% reduction in freezing during the first block of the day. Extinction training was repeated for maximum seven days.

#### Renewal and Remote Test

Twenty-four hours or 25 days after achieving the remission, animals were exposed to context D (Hybrid context) as a renewal or remote test, respectively. In each test, rats were exposed to a block of five CS+ presentations after 2 min of contextual habituation. Time intervals between tones were randomized (30-40s) during the session.

#### Immediate Footshock

To promote fear recovery, animals were placed in a neutral environment outside the conditioning box and received an unconditioned foot shock after 30 s contextual exposition, with the same intensity used during fear conditioning. The animals were returned to their home cage 30 s following the foot shock.

#### Reinstatement Test

Animals were submitted to a reinstatement test in context D twenty-four hours after the immediate footshock. Rats were exposed to a block of 5 CS+ presentations after 2 min of contextual habituation. Time intervals between tones were randomized during the session.

#### Behavioral Assessment

Freezing behavior was used as a memory index in the fear conditioning task. Freezing was analyzed off-line using Solomon software (SOLOMON CODER, (© András Péter, Budapest, Hungary), for behavioral coding by an experienced observer that was blinded to the experimental group. Freezing was defined as the absence of all movements, except those related to breathing, while the animal was alert and awake.

### Conditioned Place Preference

The conditioning box consisted of three chambers, two for the conditioning session having the same dimensions (24 × 40 × 50 cm), and the other serving as a central/start chamber (10 × 40 × 50 cm). Each chamber was employed with contextual cues and floor texture to distinguish them.

Conditioned place preference test consisted of three phases: pre-conditioning (day 1), conditioning (days 2–6) and test (day 7). The pre-conditioning session (15-min) was intended to reduce novelty and determine initial preferences for any of the two chambers by assessing the time spent in each compartment. Conditioning always took place in the initially less preferred chamber. Conditioning sessions were performed during the following five days. Animals underwent two conditioning sessions each day with 6–8-h interval between sessions. In one session, animals were placed in the initially less preferred compartment and received MFB stimulation (duration: 20 min, same intensity as used during fear conditioning experiments). During the other session the animals were placed in the opposite compartment without stimulation. The order of the sessions was randomized between animals and days. A 15-min place preference test was conducted in the absence of stimulation 24-h after the last conditioning day. The video of the animal behavior was recorded and analyzed off-line using the ANY-Maze (Stoelting, Wood Dale, IL, USA) video tracking software.

### T-Maze Task

Animals on food restriction (no less than 85% of their baseline weight) were habituated to the T-maze during 5 days before the training. The T-maze was constructed from black acrylic, with 80 cm long and 30 cm wide alleys and 40 cm high walls. Two removable doors closed the side alleys. During training, a light cue indicated the correct arm to receive a reward (froot-loops pellet). A total of 20 trials per day were performed until achieving 80% of correct choice. A removable door in the central arm was used to confine the animal at the starting point during cue presentation. After 3 min, the alley was removed, and the animal allowed to run in the maze. After arm selection, the alley was closed and the animal remains additional 3 min in the maze before next trial. Afterwards, fear conditioning, extinction and stimulation sessions started. Animals were tested in the T-maze after the extinction sessions to verify any disruption of the consolidated spatial memory. Extinction and stimulation sessions and T-maze tests were separated by five hours and the order of the behavioral tasks were randomized each day.

### Histology

Following the termination of the experiments, animals were deeply anesthetized with 1.5 g/kg urethane (i.p.) and the recording sites of each electrode were lesioned with 100 μA anodal direct current for 10 s (Supplemental Figure 1C). Then, the animals were transcardially perfused with 0.9% saline solution followed by 4% paraformaldehyde solution and 0.2% picric acid in 0.1 M phosphate buffer saline. After postfixation overnight, 50-μm thick coronal sections were prepared with a microtome (VT1000S, Leica), stained with 1 μg/ml DAPI in distilled water (D8417; Sigma-Aldrich), coverslipped and examined using a Zeiss LSM880 scanning confocal microscope (Carl Zeiss) for histological verification of the recording electrode and cannulae locations (Figure 2B and Supplemental Figure 2).

### Statistical analysis

Statistical analyses were performed using GraphPad Prism 8 software. Significance was set at p < 0.05. Data were analyzed using two-tailed Mann–Whitney U test, Kruskal–Wallis test or Mixed-Effect Analysis followed by Dunn’s post hoc or Bonferroni’s multiple comparisons test. Data are expressed as median ± IQR. For better readability, detailed statistical results and descriptive statistics are shown in Supplemental Table 1.

## ACKNOWLEDGMENTS

We thank Laura Herrera and Johanna Duran for technical assistance. This work was supported by the Momentum program II of the Hungarian Academy of Sciences (AB), EFOP-3.6.1-16-2016-00008 (AB), EFOP 3.6.6-VEKOP-16-2017-00009 (AB), and KKP133871/KKP20 grants of the National

Research, Development and Innovation Office, Hungary (AB), the 20391-3/2018/FEKUSTRAT of the Ministry of Human Capacities, Hungary, and the EU Horizon 2020 Research and Innovation Program (No. 739593—HCEMM to AB), Hungarian Scientific Research Fund (Grants NN125601 and FK123831 to MLL), the Hungarian Brain Research Program (grant KTIA_NAP_13-2-2014-0014 to MLL), UNKP-20-5 New National Excellence Program of the Ministry for Innovation and Technology from the source of the National Research, Development and Innovation Fund (MLL), Premium Postdoctoral Research Program of the Hungarian Academy of Sciences (RS). MLL was a grantee of the János Bolyai Fellowship.

## AUTHOR CONTRIBUTIONS

R.O.S., L.K.P. and A.B. conceived the project.

R.O.S., L.K.P., G.K., Y.T. and A.B. developed methodology.

R.O.S., L.K.P., L.B. and A.P. performed the experiments and analyzed data.

R.O.S., L.K.P., M.L.L., O.D., G.B. and A.B. wrote the manuscript.

O.D., G.B. advised the project.

A.B. supervised the project.

