## Supplemental Figure for "Closed-loop brain stimulation to reduce pathologic fear"

**Author affiliations:**

#### Supplemental Figure 1a

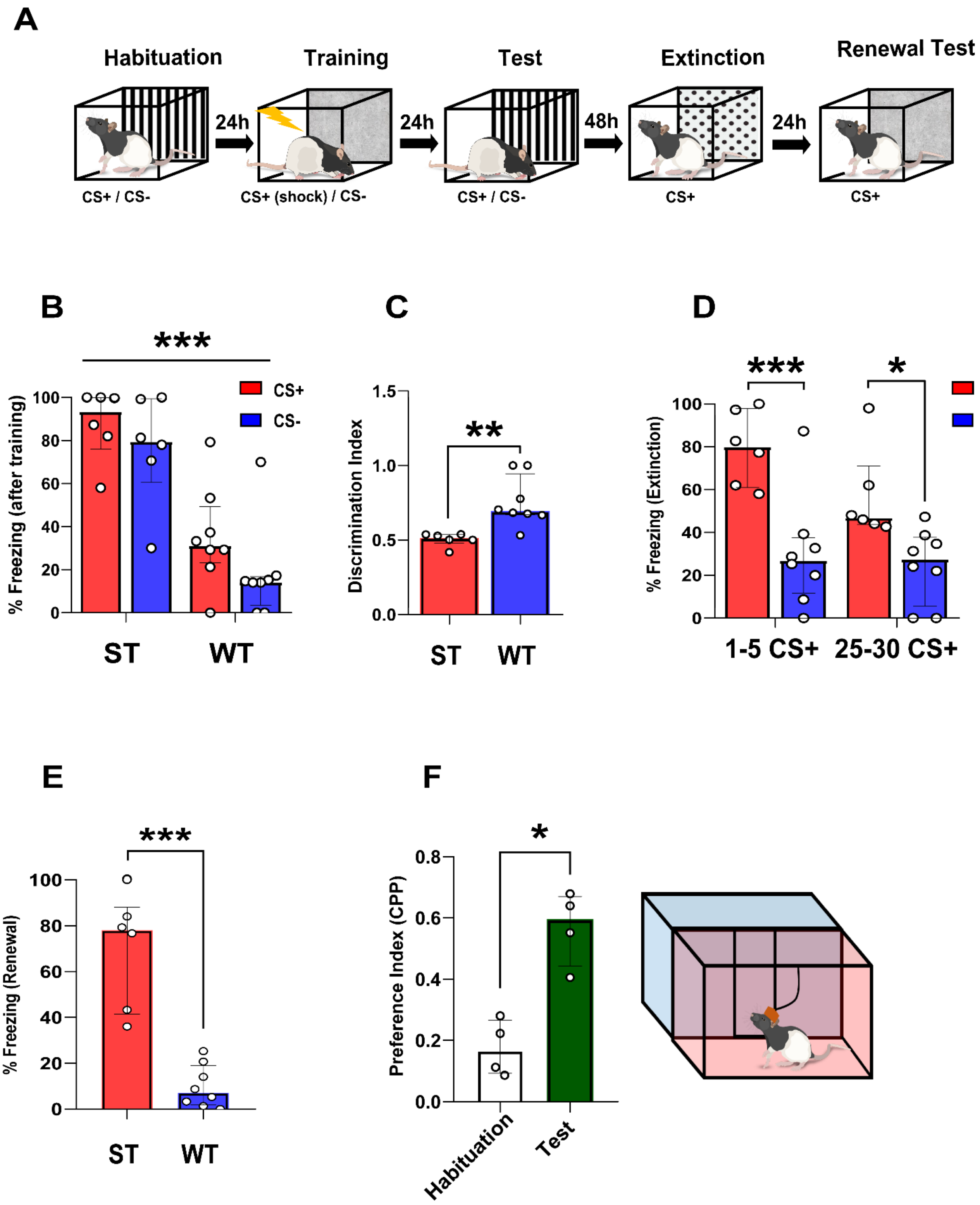

**Supplemental Figure 1a. Strong fear conditioning induces a PTSD-like phenotype.** (**A**) Schematics of the experimental design. (B) Animals submitted to strong training (ST) during fear conditioning (5x1mA) showed robust fear reaction compared to a week training (WT) (5x0.4mA) either CS+ and CS- test. However, there is no significant differences within the same group regarding fear reactions to CS+ or CS- (Group factor (strong vs week) $F (1, 12) = 22.68$, time factor (CS+ vs CS-) $F (1, 12) = 23.88$, group x time interaction $F (1, 12) = 1.023$, $*p < 0.001$ in in Bonferroni's multiple comparisons after Mixed-effects test). (**C**)**.** Fear generalization analysis using a discrimination index showed that strong training induces poor discrimination between CS+ and CS- compared to a week training ($U = 2.00$, $p = 0.0027$; $**p < 0.01$ in Mann–Whitney test). **(D).** During extinction, animals in the strong training group expressed high fear responses during the initial and last 5 blocks of CS+ compared to the week training. Moreover, there is no significant decrease of freezing levels in both groups across the extinction (0-5min to 25-30min) (Group factor (strong vs week) $F (1, 14) = 3.061$, $p=0.1021$; time factor (0-5min to 25-30min) $F (1, 10) = 25.35$, $p < 0.001$; group x time interaction $F (1, 10) = 1.144$, $p= 0.3100$; $***p < 0.001$ in in Bonferroni's multiple comparisons after Mixed-effects test) (Strong training n=6; Weak training n=8). **(E).** Animas exposed to renewal test in a hybrid context and submitted to strong fear conditioning expresses high fear reactions compared to animals trained with a week training ($U = 0.00$, $p = 0.0007$; $***p < 0.001$ in Mann–Whitney test) ST, n=6; WT, n=8. **(F).** In order to assess the rewarding properties of the MFB stimulation, animals were submitted to conditioning place preference (see Material and Methods). Animals expressed a significant preference for the chamber paired with the MFB stimulation during the test compared to the habituation session ($U = 0.00$, $p = 0.0286$; $*p < 0.005$ in Mann–Whitney test, n=4). * = $p< 0.05$, ** = $p< 0.01$, *** = $p< 0.001$.

#### Supplemental Figure 1b

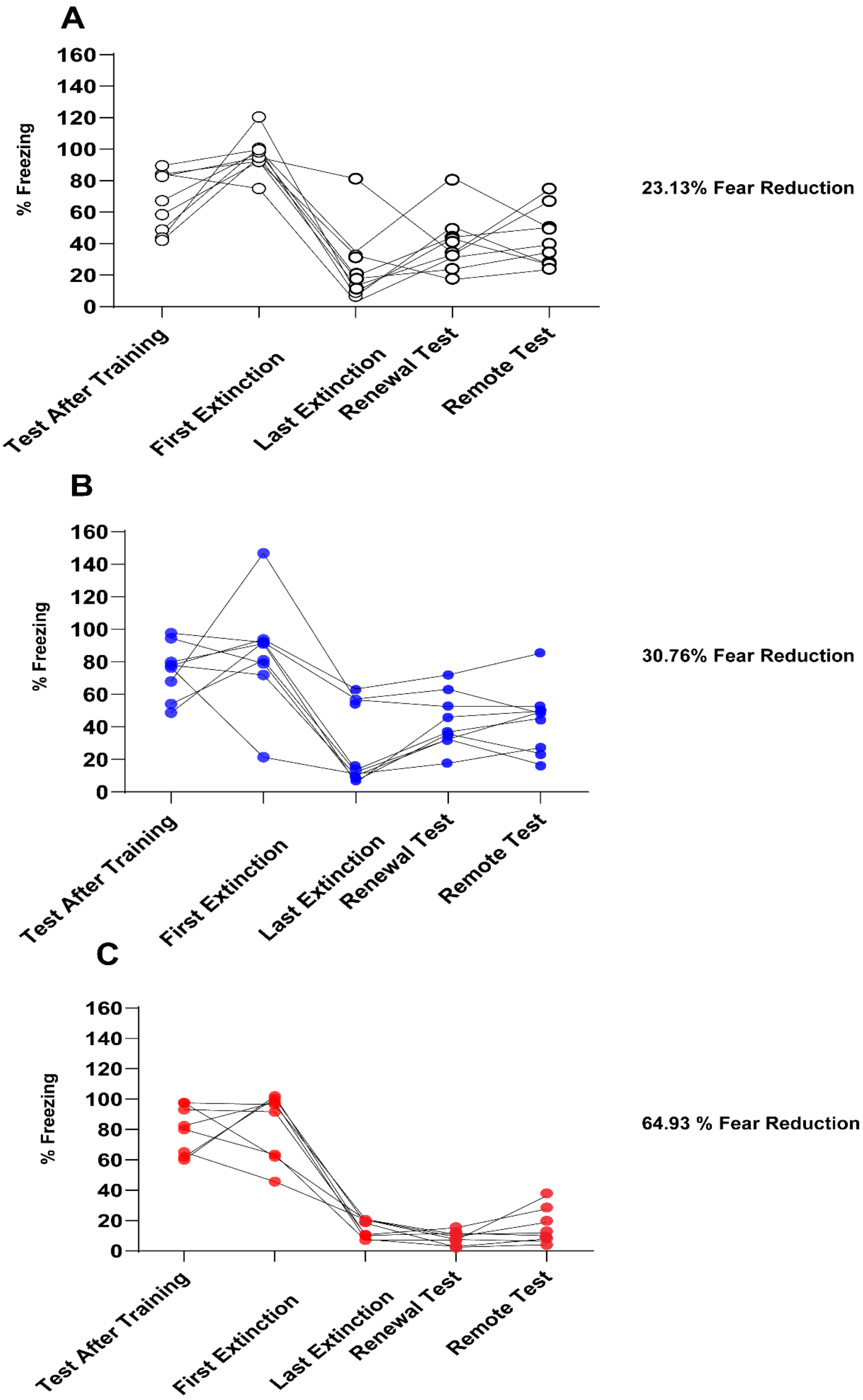

**Supplemental Figure 1b. Individual fear expression across sessions.** Graph displaying individual performance across the experiment. Percentage (right to each graph) shows fear reduction from a $\Delta$ freezing analysis (see Figure 1 I) comparing test to CS+ and remote test (25 days after extinction). Non-stimulated (23.13%, n=9), Open-Loop (30.76%, n=9), Closed-loop (64.93%, n=8).

### Supplemental Figure 1c

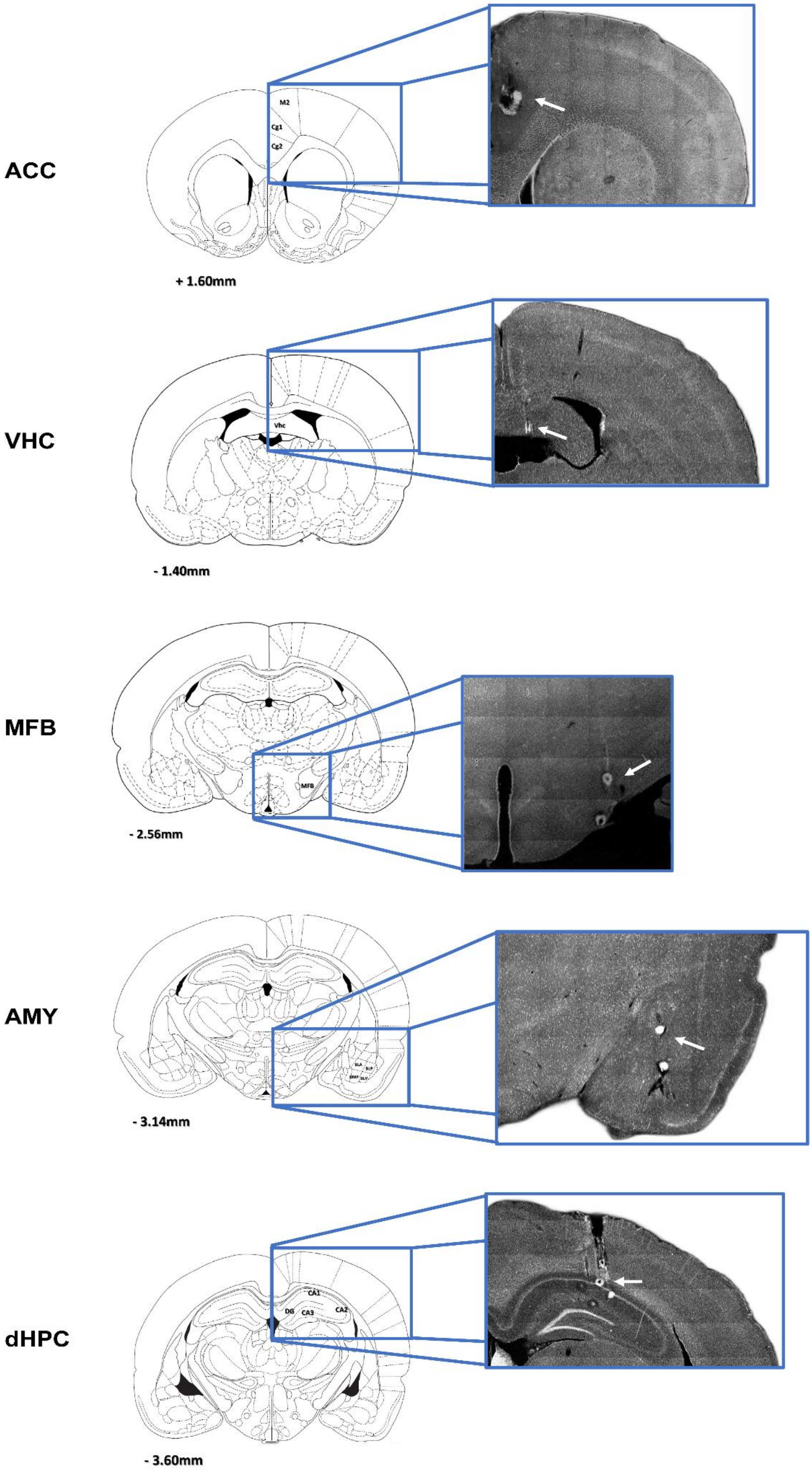

**Supplemental Figure 1c.** Histological verification of electrode placements. A coronal section of the anterior cingulate cortex (ACC), ventral hippocampal commissure (VHC), medial forebrain bundle (MFB), Amygdala (Amy) and dorsal hippocampus (dHPC) stained with DAPI is shown. recording sites on each shank were lesioned (white dots) after the end of the experiment by applying 100 μA anodal direct current for 10 s via electrode tips.

#### Supplemental Figure 2

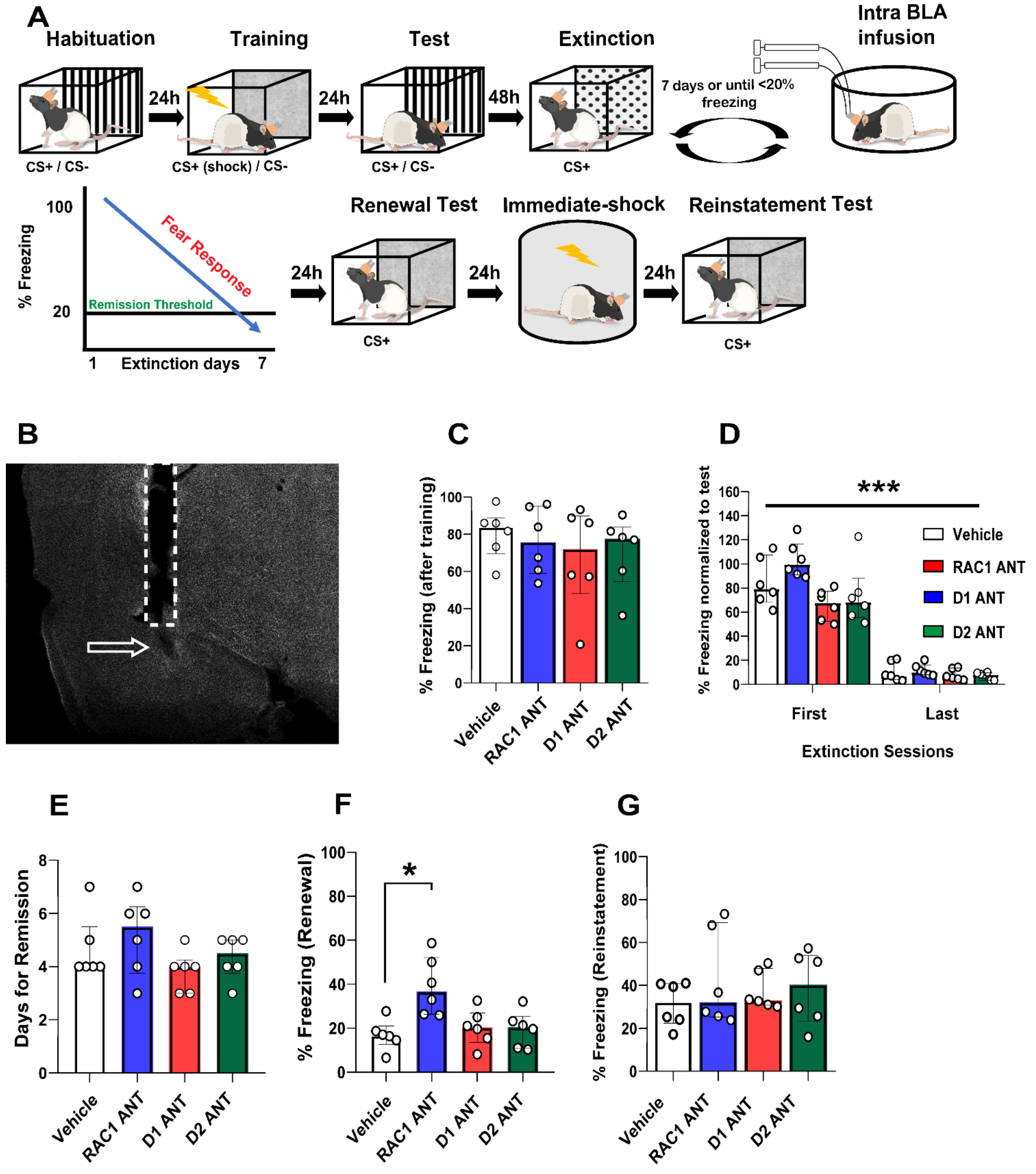

**Supplemental Figure 2. Pharmacological evaluation of NSC2376, SCH23390 and Sulpiride in absence of closed-loop stimulation.** **(A)** Schematics of the experimental design. **(B)** Representative histological verification of a cannula placement. **(C)** During the test to the CS+, there are no significant differences between the groups, showing that all groups are able to acquire fear conditioning in a similar fashion. **(D)** No differences were found in the fear expression between groups during the first 5 CS+ block from first and last extinction day. However, there is a significant decrease in fear expression over time (Group factor F (2.181, 21.81) = 5.137, p= 0.0131; time factor (first extinction vs last extinction) F (1, 10) = 318.9, p < 0.001; group x time interaction F (3, 30) = 3.131, p=0.0401; ***p < 0.001 in Bonferroni's multiple comparisons after Mixed-effects test). **(E).** The drugs do not have effect itself over the extinction criterion **(F).** However, during the renewal, NSC2376 seems to increase freezing behavior only compared to control animals infused with saline (H = 10.86, p = 0.0125; *p < 0.05 in Dunn's multiple comparisons after Kruskal–Wallis test). **()** No differences were detected during the reinstatement test. Control (Saline) n=6; NSC2376 n=6; SCH23390 n=6; Sulpiride n=6. * = p< 0.05, *** = p< 0.001.

#### Supplemental Figure 3

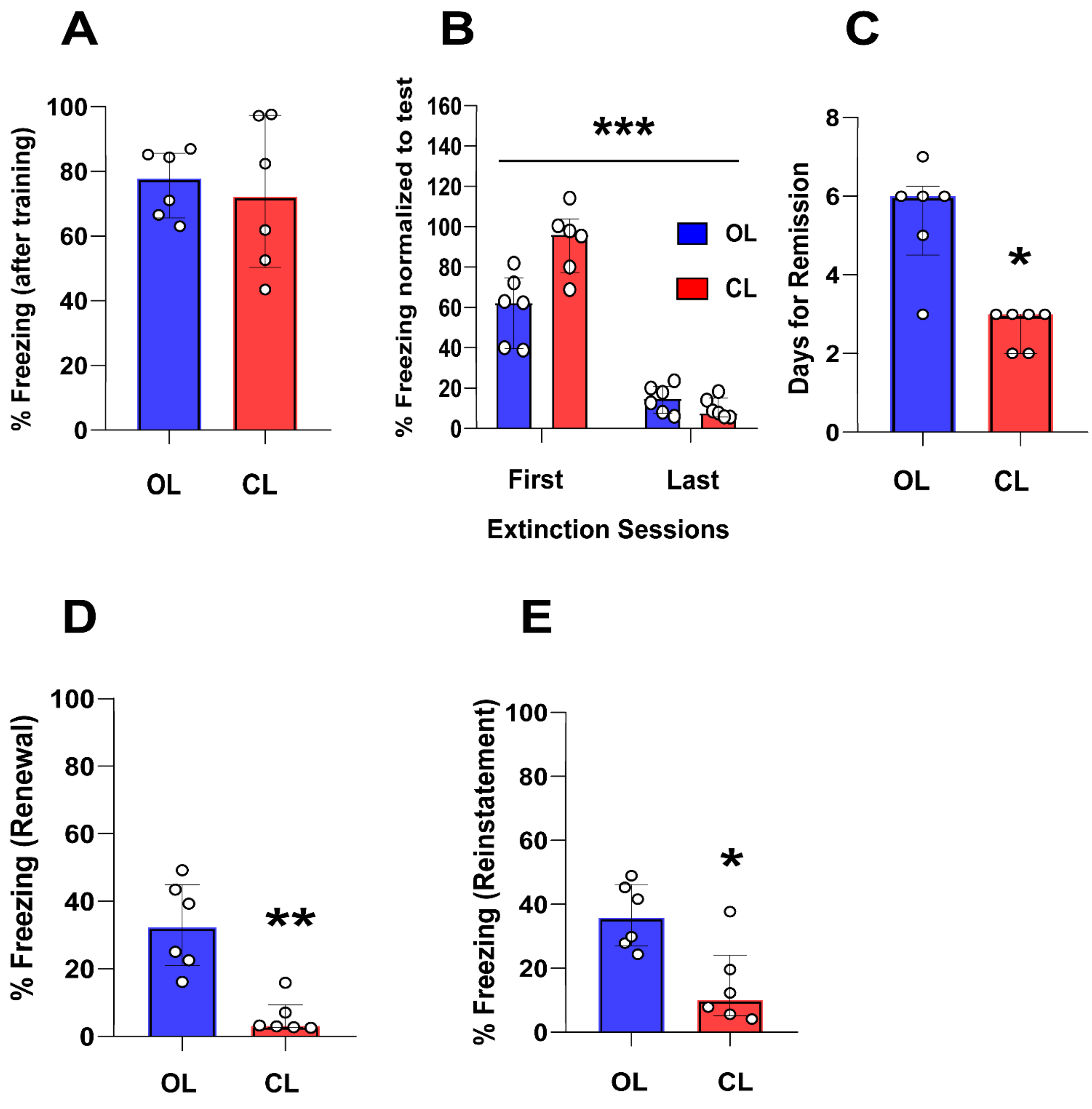

**Supplemental Figure 3. Extinction enhancement induced by closed-loop neuromodulation is not affected by previously consolidated memories.** Data from animals submitted to concomitant T-maze task and closed-loop neuromodulation for extinction enhancement. **A.** During the test to the CS+, there are no significant differences between the groups showing that all groups are able to acquire the fear conditioning. **B.** No differences were found in the fear expression between groups during the first 5 CS+ block from first and last extinction day. However, there is a significant decrease in fear expression over time (Group factor $F (1, 10) = 5.473$, $p= 0.0414$; time factor (first extinction vs last extinction) $F (1, 10) = 261.9$, $p < 0.0001$; group x time interaction $F (1, 10) = 22.90$, $p=0.0007$; ***$p < 0.001$ in Bonferroni's multiple comparisons after Mixed-effects test). **C.** Animals exposed to closed-loop stimulation required less extinction sessions to achieve the remission criterion compared to the open-loop group ($U = 2.00$, $p = 0.0108$; *$p < 0.05$ in Mann–Whitney test). **D.** The enhancement was also expressed during renewal test since closed-loop neuromodulation induce lower fear expression ($U = 0.00$, $p = 0.0022$; **$p < 0.01$ in Mann–Whitney test). **E.** During the reinstatement test, closed-loop prevents the fear recovery induced by an immediate foot-shock procedure ($U = 3.00$, $p = 0.0152$; *$p < 0.05$ in Mann–Whitney test). Open-loop (OL) n=6, Closed-loop (CL) n=6. * = $p< 0.05$, ** = $p< 0.01$, *** = $p< 0.001$.
