## Supplemental Table for "Closed-loop brain stimulation to reduce pathologic fear"

| Figure | Panel Title | Conclusion | Supporting Data & Statistics | | | | |
|---|---|---|---|---|---|---|---|
| 1D | Test After Training | No difference in fear expression in response to the CS+ following training | Number of animals | 9 | 9 | 8 | |
| | | | | **Non-stimulated** | **Open-Loop** | **Closed-Loop** | |
| | | | 25% Percentile | 46,13 | 60,93 | 62,67 | |
| | | | Median | 67,33 | 77,73 | 81,27 | |
| | | | 75% Percentile | 84,13 | 87,13 | 96,37 | |
| | | | Kruskal-Wallis test | | | | |
| | | | P value | 0,4195 | | | |
| | | | Number of groups | 3 | | | |
| | | | Kruskal-Wallis statistic | 1,737 | | | |
| 1E | Transition to Extinction | All groups are able to decrease the fear response over time | 25% Percentile | 14,69 | 23,65 | | |
| | | | Median | 46,84 | 56,82 | | |
| | | | 75% Percentile | 79 | 89,99 | | |
| | | | Mixed-Effect Analysis | P value | P value summary | F (DFn, DFd) | |
| | | | Time Factor (1-5 vs 15-20 tones) | <0.0001 | **** | F (1, 23) = 164.2 | |
| | | | Group Factor | 0,3229 | ns | F (2, 23) = 1.188 | |
| | | | Group Factor x Time Factor | 0,4859 | ns | F (2, 23) = 0.7450 | |
| | | | Bonferroni's multiple comparisons test | Predicted (LS) mean diff. | 95.00% CI of diff. | Summary | Adjusted P Value |
| | | | 1-5 vs 15-20 tones | | | | |
| | | | Non-stimulated | 74,43 | 51.46 to 97.41 | **** | <0.0001 |
| | | | Open Loop | 59,13 | 36.16 to 82.10 | **** | <0.0001 |
| | | | Closed Loop | 67,97 | 43.60 to 92.33 | **** | <0.0001 |
| 1F | Days for Crossing the Threshold | Animals exposed to closed-loop stimulation required less extinction sessions to achieve the remission criterion | 25% Percentile | 4 | 5 | 2 | |
| | | | Median | 5 | 6 | 2 | |
| | | | 75% Percentile | 7 | 7 | 4 | |
| | | | Kruskal-Wallis test | | | | |
| | | | P value | 0,0011 | | | |
| | | | Kruskal-Wallis statistic | 13,6 | | | |
| | | | Dunn's multiple comparisons test | Mean rank diff. | Summary | Adjusted P Value | |
| | | | Non-Stimulated vs. Open Loop | -2,167 | ns | >0.9999 | |
| | | | Non-Stimulated vs. Closed Loop | 10,56 | * | 0,0118 | |
| | | | Open Loop vs. Closed Loop | 12,73 | ** | 0,0015 | |
| | | | Test details | Mean rank 1 | Mean rank 2 | Mean rank diff. | |
| | | | Non-Stimulated vs. Open Loop | 16 | 18,17 | -2,167 | |
| | | | Non-Stimulated vs. Closed Loop | 16 | 5,438 | 10,56 | |
| | | | Open Loop vs. Closed Loop | 18,17 | 5,438 | 12,73 | |
| 1G | Renewal Test | Closed-loop neuromodulation induced lower fear expression during the renewal test | 25% Percentile | 27,33 | 32,33 | 3,967 | |
| | | | Median | 34 | 36,8 | 8,373 | |
| | | | 75% Percentile | 47,33 | 57,73 | 11,4 | |
| | | | Kruskal-Wallis test | | | | |
| | | | P value | 0,0003 | | | |
| | | | Kruskal-Wallis statistic | 16,21 | | | |
| | | | Dunn's multiple comparisons test | Mean rank diff. | Summary | Adjusted P Value | |
| | | | Non-stimulated vs. Open Loop | -1,667 | ns | >0.9999 | |
| | | | Non-stimulated vs. Closed Loop | 12,17 | ** | 0,0032 | |
| | | | Open Loop vs. Closed Loop | 13,83 | *** | 0,0006 | |
| 1H | Remote Test | Closed-loop neuromodulation prevented spontaneous fear recovery 25 days after extinction | 25% Percentile | 26,8 | 25,47 | 6,667 | |
| | | | Median | 39,33 | 48,27 | 10,93 | |
| | | | 75% Percentile | 58,53 | 51 | 25,53 | |
| | | | Kruskal-Wallis test | | | | |
| | | | P value | 0,0056 | | | |
| | | | Kruskal-Wallis statistic | 10,38 | | | |
| | | | Dunn's multiple comparisons test | Mean rank diff. | Summary | Adjusted P Value | |
| | | | Non-Stimulated vs. Open Loop | 0,1111 | ns | >0.9999 | |
| | | | Non-Stimulated vs. Closed Loop | 10,53 | * | 0,0138 | |
| | | | Open Loop vs. Closed Loop | 10,42 | * | 0,0152 | |
| 1I | Remote Test - Test After Training | Closed-loop stimulated animals had stronger fear reduction over time | 25% Percentile | -37,2 | -49,27 | -79,27 | |
| | | | Median | -17,2 | -30,27 | -62,07 | |
| | | | 75% Percentile | -12,67 | -9,467 | -51,93 | |
| | | | Kruskal-Wallis test | | | | |
| | | | P value | 0,0015 | | | |
| | | | Kruskal-Wallis statistic | 13,06 | | | |
| | | | Dunn's multiple comparisons test | Mean rank diff. | Summary | Adjusted P Value | |
| | | | Non-Stimulated vs. Open Loop | 2,333 | ns | >0.9999 | |
| | | | Non-Stimulated vs. Closed Loop | 12,72 | ** | 0,0019 | |
| | | | Open Loop vs. Closed Loop | 10,39 | * | 0,0156 | |
| 2B | Test After Training | No difference in fear expression in response to the CS+ following training | Number of animals | 5 | 5 | | |
| | | | | **Non-stimulated** | **Closed-Loop** | | |
| | | | 25% Percentile | 77,67 | 84,87 | | |
| | | | Median | 90,93 | 86,27 | | |
| | | | 75% Percentile | 95,93 | 91,53 | | |
| | | | P value | 0,3095 | | | |
| | | | Sum of ranks in column A,B | 33 , 22 | | | |
| | | | Mann-Whitney U | 7 | | | |
| 2C | Renewal Test | Closed-loop neuromodulation without extinction fails to reduce fear expression during the renewal test | 25% Percentile | 67,4 | 79,53 | | |
| | | | Median | 83,47 | 88,27 | | |
| | | | 75% Percentile | 86,53 | 94,33 | | |
| | | | P value | 0,2222 | | | |
| | | | Sum of ranks in column A,B | 21 , 34 | | | |
| | | | Mann-Whitney U | 6 | | | |
| 2E-G | T-maze performance | No differences were detected between groups during the T-maze and there is no decay of the performance over time within groups | Number of Animals | 5 | 5 | | |
| | | | | **Open-Loop** | **Closed-loop** | | |
| | | | Mean | 81,17 | 83,83 | | |
| | | | Std. Deviation | 7,897 | 5,29 | | |
| | | | Std. Error of Mean | 3,532 | 2,366 | | |
| | | | Unpaired t test | P value | Mean of Open Loop | Mean of Closed Loop | Difference |
| | | | Before CFC | 0,270454 | 80 | 89,17 | -9,167 |
| | | | After CFC | 0,059954 | 69,17 | 85 | -15,83 |
| | | | First Extinction | 0,614714 | 80 | 75,83 | 4,167 |
| | | | Last Extinction | 0,421528 | 88,33 | 81,67 | 6,667 |
| | | | Renewal Test | 0,919713 | 88,33 | 87,5 | 0,8333 |
| | | | Number of Animals | 6 | 5 | 5 | 5 |
| | | | | **Closed Loop (CL)** | **CL + NSC 23766** | **CL + SCH 23390** | **CL + SULPIRIDE** |
| | | | 25% Percentile | 53,27 | 62,27 | 57,8 | 86,07 |
| | | | Median | 66,27 | 84,27 | 61,6 | 94,8 |
| | | | 75% Percentile | 80,44 | 89,93 | 93,4 | 98 |
| | | | Kruskal-Wallis test | | | | |
| | | | P value | 0,1429 | | | |
| | | | Kruskal-Wallis statistic | 5,43 | | | |
| | | | Dunn's multiple comparisons test | Mean rank diff. | Summary | Adjusted P Value | |

| Figure | Panel Title | Conclusion | Supporting Data & Statistics | | | | |
|---|---|---|---|---|---|---|---|
| 3C | Test After Training | No difference in fear expression in response to the CS+ following training | Closed Loop (CL) vs. CL + NSC 23766 | -3,333 | ns | >0.9999 | |
| | | | Closed Loop (CL) vs. CL + SCH 23390 | -2,133 | ns | >0.9999 | |
| | | | Closed Loop (CL) vs. CL + SULPIRIDE | -8,533 | ns | 0,1388 | |
| | | | CL + NSC 23766 vs. CL + SCH 23390 | 1,2 | ns | >0.9999 | |
| | | | CL + NSC 23766 vs. CL + SULPIRIDE | -5,2 | ns | >0.9999 | |
| | | | CL + SCH 23390 vs. CL + SULPIRIDE | -6,4 | ns | 0,6175 | |
| | | | Test details | Mean rank 1 | Mean rank 2 | Mean rank diff. | |
| | | | Closed Loop (CL) vs. CL + NSC 23766 | 7,667 | 11 | -3,333 | |
| | | | Closed Loop (CL) vs. CL + SCH 23390 | 7,667 | 9,8 | -2,133 | |
| | | | Closed Loop (CL) vs. CL + SULPIRIDE | 7,667 | 16,2 | -8,533 | |
| | | | CL + NSC 23766 vs. CL + SCH 23390 | 11 | 9,8 | 1,2 | |
| | | | CL + NSC 23766 vs. CL + SULPIRIDE | 11 | 16,2 | -5,2 | |
| | | | CL + SCH 23390 vs. CL + SULPIRIDE | 9,8 | 16,2 | -6,4 | |
| 3D | Transition to Extinction | All groups are able to decrease the fear response over time | 25% Percentile | 12,87 | 19,98 | 15,23 | 10,91 |
| | | | Median | 45,66 | 52,17 | 44,2 | 46,84 |
| | | | 75% Percentile | 78,45 | 84,36 | 73,16 | 82,78 |
| | | | Mixed-Effect Analysis | P value | P value summary | F (DFn, DFd) | |
| | | | Time Factor (1-5 vs 15-20 tones) | <0.0001 | **** | F (1, 34) = 175.1 | |
| | | | Group Factor | 0,694 | ns | F (3, 34) = 0.4864 | |
| | | | Group Factor x Time Factor | 0,8072 | ns | F (3, 34) = 0.3251 | |
| | | | Bonferroni's multiple comparisons test | Predicted (LS) mean diff. | 95.00% CI of diff. | Summary | Adjusted P Value |
| | | | 1-5 vs 15-20 tones | | | | |
| | | | Closed Loop (CL) | 65,58 | 41.44 to 89.72 | **** | <0.0001 |
| | | | CL + NSC 23766 | 64,39 | 37.94 to 90.83 | **** | <0.0001 |
| | | | CL + SCH 23390 | 57,93 | 31.49 to 84.38 | **** | <0.0001 |
| | | | CL + SULPIRIDE | 71,88 | 45.43 to 98.32 | **** | <0.0001 |
| 3E | Days for Crossing the Threshold | NSC2376 and sulpiride injected animals required more extinction sessions to achieve the extinction criterion | 25% Percentile | 2 | 5 | 2 | 4 |
| | | | Median | 2,5 | 5 | 2 | 5 |
| | | | 75% Percentile | 3 | 6 | 2,5 | 6,5 |
| | | | Kruskal-Wallis test | | | | |
| | | | P value | 0,0011 | | | |
| | | | Kruskal-Wallis statistic | 16,16 | | | |
| | | | Dunn's multiple comparisons test | Mean rank diff. | Summary | Adjusted P Value | |
| | | | Closed Loop (CL) vs. CL + NSC 23766 | -10,15 | * | 0,0324 | |
| | | | Closed Loop (CL) vs. CL + SCH 23390 | 1,65 | ns | >0.9999 | |
| | | | Closed Loop (CL) vs. CL + SULPIRIDE | -9,35 | ns | 0,0623 | |
| | | | CL + NSC 23766 vs. CL + SCH 23390 | 11,8 | * | 0,0117 | |
| | | | CL + NSC 23766 vs. CL + SULPIRIDE | 0,8 | ns | >0.9999 | |
| | | | CL + SCH 23390 vs. CL + SULPIRIDE | -11 | * | 0,0234 | |
| | | | Test details | Mean rank 1 | Mean rank 2 | Mean rank diff. | |
| | | | Closed Loop (CL) vs. CL + NSC 23766 | 6,75 | 16,9 | -10,15 | |
| | | | Closed Loop (CL) vs. CL + SCH 23390 | 6,75 | 5,1 | 1,65 | |
| | | | Closed Loop (CL) vs. CL + SULPIRIDE | 6,75 | 16,1 | -9,35 | |
| | | | CL + NSC 23766 vs. CL + SCH 23390 | 16,9 | 5,1 | 11,8 | |
| | | | CL + NSC 23766 vs. CL + SULPIRIDE | 16,9 | 16,1 | 0,8 | |
| | | | CL + SCH 23390 vs. CL + SULPIRIDE | 5,1 | 16,1 | -11 | |
| 3F | Renewal Test | Sulpiride suppress the extinction enhancement induced by closed-loop neuromodulation during renewal | 25% Percentile | 2,533 | 21 | 1 | 28 |
| | | | Median | 3,198 | 28,13 | 4,8 | 38,93 |
| | | | 75% Percentile | 7,067 | 38,2 | 8 | 52,2 |
| | | | Kruskal-Wallis test | | | | |
| | | | P value | 0,002 | | | |
| | | | Kruskal-Wallis statistic | 14,84 | | | |
| | | | Dunn's multiple comparisons test | Mean rank diff. | Summary | Adjusted P Value | |
| | | | Closed Loop (CL) vs. CL + NSC 23766 | -9,2 | ns | 0,086 | |
| | | | Closed Loop (CL) vs. CL + SCH 23390 | -0,2 | ns | >0.9999 | |
| | | | Closed Loop (CL) vs. CL + SULPIRIDE | -11,6 | * | 0,0121 | |
| | | | CL + NSC 23766 vs. CL + SCH 23390 | 9 | ns | 0,1309 | |
| | | | CL + NSC 23766 vs. CL + SULPIRIDE | -2,4 | ns | >0.9999 | |
| | | | CL + SCH 23390 vs. CL + SULPIRIDE | -11,4 | * | 0,022 | |
| | | | Test details | Mean rank 1 | Mean rank 2 | Mean rank diff. | |
| | | | Closed Loop (CL) vs. CL + NSC 23766 | 6 | 15,2 | -9,2 | |
| | | | Closed Loop (CL) vs. CL + SCH 23390 | 6 | 6,2 | -0,2 | |
| | | | Closed Loop (CL) vs. CL + SULPIRIDE | 6 | 17,6 | -11,6 | |
| | | | CL + NSC 23766 vs. CL + SCH 23390 | 15,2 | 6,2 | 9 | |
| | | | CL + NSC 23766 vs. CL + SULPIRIDE | 15,2 | 17,6 | -2,4 | |
| | | | CL + SCH 23390 vs. CL + SULPIRIDE | 6,2 | 17,6 | -11,4 | |
| 3G | Reinstatement Test | Animals treated with NSC2376 and sulpiride are more prone to fear recovery compared to the other groups | 25% Percentile | 5,369 | 17,87 | 4,533 | 16,73 |
| | | | Median | 6,667 | 27,07 | 11,2 | 28,4 |
| | | | 75% Percentile | 10,97 | 43,8 | 15,2 | 30,93 |
| | | | Kruskal-Wallis test | | | | |
| | | | P value | 0,0057 | | | |
| | | | Kruskal-Wallis statistic | 12,55 | | | |
| | | | Dunn's multiple comparisons test | Mean rank diff. | Summary | Adjusted P Value | |
| | | | Closed Loop (CL) vs. CL + NSC 23766 | -9,967 | * | 0,0478 | |
| | | | Closed Loop (CL) vs. CL + SCH 23390 | -1,367 | ns | >0.9999 | |
| | | | Closed Loop (CL) vs. CL + SULPIRIDE | -10,37 | * | 0,0347 | |
| | | | CL + NSC 23766 vs. CL + SCH 23390 | 8,6 | ns | 0,1702 | |
| | | | CL + NSC 23766 vs. CL + SULPIRIDE | -0,4 | ns | >0.9999 | |
| | | | CL + SCH 23390 vs. CL + SULPIRIDE | -9 | ns | 0,1307 | |
| | | | Test details | Mean rank 1 | Mean rank 2 | Mean rank diff. | |
| | | | Closed Loop (CL) vs. CL + NSC 23766 | 5,833 | 15,8 | -9,967 | |
| | | | Closed Loop (CL) vs. CL + SCH 23390 | 5,833 | 7,2 | -1,367 | |
| | | | Closed Loop (CL) vs. CL + SULPIRIDE | 5,833 | 16,2 | -10,37 | |
| | | | CL + NSC 23766 vs. CL + SCH 23390 | 15,8 | 7,2 | 8,6 | |
| | | | CL + NSC 23766 vs. CL + SULPIRIDE | 15,8 | 16,2 | -0,4 | |
| | | | CL + SCH 23390 vs. CL + SULPIRIDE | 7,2 | 16,2 | -9 | |
| F4C | Test After Training | No difference in fear expression in response to the CS+ following training | Number of Animals | 9 | 5 | | |
| | | | | Open-Loop | Closed-loop | | |
| | | | 25% Percentile | 43,93 | 51,47 | | |
| | | | Median | 61,2 | 79,2 | | |
| | | | 75% Percentile | 77,27 | 92,67 | | |
| | | | Mann Whitney test | | | | |
| | | | P value | 0,2977 | | | |
| | | | Sum of ranks in column A,B | 59 , 46 | | | |
| | | | Mann-Whitney U | 14 | | | |
| F4D | Transition to Extinction | All groups are able to decrease the fear response over time | 25% Percentile | 14,69 | 23,65 | | |
| | | | Median | 46,84 | 56,82 | | |
| | | | 75% Percentile | 79 | 89,99 | | |
| | | | Bonferroni's multiple comparisons test | Summary | Adjusted P Value | | |

| Figure | Panel Title | Conclusion | Supporting Data & Statistics | | | | |
|---|---|---|---|---|---|---|---|
| | | | 1-5 vs 15-20 tones | | | | |
| | | | Open Loop | **** | <0.0001 | | |
| | | | Closed Loop | *** | 0,0005 | | |
| F4E | Days for Crossing the Threshold | SWR disrupted animals require more days to achieve the extinction criterion | 25% Percentile | 2 | 3 | | |
| | | | Median | 3 | 4 | | |
| | | | 75% Percentile | 3 | 6 | | |
| | | | Mann Whitney test | | | | |
| | | | P value | 0,028 | | | |
| | | | Sum of ranks in column A,B | 51 , 54 | | | |
| | | | Mann-Whitney U | 6 | | | |
| F4F | Renewal Test | SWR disrupted animals show high fear expression during renewal | 25% Percentile | 5,2 | 23,6 | | |
| | | | Median | 10,67 | 43,47 | | |
| | | | 75% Percentile | 20,13 | 60,93 | | |
| | | | Mann Whitney test | | | | |
| | | | P value | 0,012 | | | |
| | | | Sum of ranks in column A,B | 49 , 56 | | | |
| | | | Mann-Whitney U | 4 | | | |
| F4G | Reinstatement Test | No difference in fear expression during reinstatement | 25% Percentile | 38,53 | 29 | | |
| | | | Median | 46,93 | 65,6 | | |
| | | | 75% Percentile | 60,27 | 88,07 | | |
| | | | Mann Whitney test | | | | |
| | | | P value | 0,7972 | | | |
| | | | Sum of ranks in column A,B | 65 , 40 | | | |
| | | | Mann-Whitney U | 20 | | | |
| S1aB | Test After Training | No difference in fear expression in response to the CS+ following training | Number of animals | 6 | 8 | | |
| | | | | CS+ | CS- | | |
| | | | 25% Percentile | 35,42 | 18,17 | | |
| | | | Median | 61,65 | 47,36 | | |
| | | | 75% Percentile | 87,89 | 76,56 | | |
| | | | Mixed-Effect Analysis | P value | P value summary | lly significant (P < 0.05)? | F (DFn, DFd) |
| | | | Training Intensity (strong vs weak) Factor | 0,0005 | *** | Yes | F (1, 12) = 22.68 |
| | | | Test (CS+ vs CS-)Time | 0,0004 | *** | Yes | F (1, 12) = 23.88 |
| | | | Training Factor x Test | 0,3317 | ns | No | F (1, 12) = 1.023 |
| | | | Bonferroni's multiple comparisons test | Predicted (LS) mean diff. | 95.00% CI of diff. | | |
| | | | CS+ - CS- | | | | |
| | | | Strong Training | 11,33 | 0.01445 to 22.65 | | |
| | | | Weak Training | 17,25 | 7.448 to 27.05 | | |
| S1aC | y Axis showing Discrmination Index | Strong training induces poor discrimination between CS+ and CS- | 25% Percentile | 0,4794 | 0,6692 | | |
| | | | Median | 0,5149 | 0,6936 | | |
| | | | 75% Percentile | 0,5373 | 0,9442 | | |
| | | | Mann Whitney test | | | | |
| | | | P value | 0,0027 | | | |
| | | | Sum of ranks in column A,B | 23 , 82 | | | |
| | | | Mann-Whitney U | 2 | | | |
| S1aD | Extinction | The strong training group expressed high fear responses during the initial and last 5 blocks of CS+ compared to the week training. Moreover, there is no significant decrease of freezing levels in both groups across the extinction (0-5min to 25-30min) | 25% Percentile | 30,25 | 24,75 | | |
| | | | Median | 54,9 | 40,76 | | |
| | | | 75% Percentile | 79,56 | 56,78 | | |
| | | | Mixed-Effect Analysis | P value | P value summary | F (DFn, DFd) | |
| | | | Training Intensity (strong vs weak) Factor | 0,0005 | *** | F (1, 12) = 21.96 | |
| | | | Test (CS+ vs CS-)Time | 0,0814 | ns | F (1, 12) = 3.619 | |
| | | | Training Factor x Test | 0,2677 | ns | F (1, 12) = 1.351 | |
| | | | Bonferroni's multiple comparisons test | Predicted (LS) mean diff. | 95.00% CI of diff. | Summary | Adjusted P Value |
| | | | Strong Training - Weak Training | | | | |
| | | | (CS+) 0-5min | 49,31 | 21.99 to 76.62 | *** | 0,0005 |
| | | | (CS+) 25-30min | 32,03 | 4.710 to 59.35 | * | 0,0197 |
| S1aE | Renewal Test | Strong fear conditioning induce high fear reactions during a renewal test | 25% Percentile | 41,5 | 1,833 | | |
| | | | Median | 78 | 7 | | |
| | | | 75% Percentile | 88 | 19 | | |
| | | | Mann Whitney test | | | | |
| | | | P value | 0,0007 | | | |
| | | | Sum of ranks in column A,B | 69 , 36 | | | |
| | | | Mann-Whitney U | 0 | | | |
| S1aF | Preference Stimulated Compartiment | Animals expressed a significant preference for the chamber paired with the MFB stimulation during the test compared to the habituation session | Number of animals | 4 | | | |
| | | | 25% Percentile | 0,09289 | 0,4423 | | |
| | | | Median | 0,1676 | 0,5957 | | |
| | | | 75% Percentile | 0,2662 | 0,6693 | | |
| | | | Mann Whitney test | | | | |
| | | | P value | 0,0286 | | | |
| | | | Sum of ranks in column A,B | 10 , 26 | | | |
| | | | Mann-Whitney U | 0 | | | |
| S2C | Test After Training | No difference in fear expression in response to the CS+ following training | Number of Animals | 6 | 6 | 6 | 6 |
| | | | | **Control** | **NSC 23766** | **SCH 23390** | **SULPIRIDE** |
| | | | 25% Percentile | 69,43 | 58,9 | 48,1 | 54,37 |
| | | | Median | 83,8 | 75,67 | 72,07 | 77,67 |
| | | | 75% Percentile | 88,9 | 95,13 | 89,9 | 83,8 |
| | | | Kruskal-Wallis test | | | | |
| | | | P value | 0,7867 | | | |
| | | | Kruskal-Wallis statistic | 1,06 | | | |
| S2D | Transition to Extinction | All groups are able to decrease the fear response over time | 25% Percentile | 10,5 | 11,95 | 8,017 | 7,227 |
| | | | Median | 47,79 | 57,77 | 37,09 | 40,66 |
| | | | 75% Percentile | 85,08 | 103,6 | 66,16 | 74,09 |
| | | | Mixed-Effect Analysis | P value | P value summary | F (DFn, DFd) | |
| | | | Time Factor (1-5 vs 15-20 tones) | <0.0001 | **** | F (1, 10) = 318.9 | |
| | | | Group Factor | 0,0131 | * | F (2.181, 21.81) = 5.137 | |
| | | | Group Factor x Time Factor | 0,0401 | * | F (3, 30) = 3.131 | |
| | | | Bonferroni's multiple comparisons test | Predicted (LS) mean diff. | 95.00% CI of diff. | Summary | Adjusted P Value |
| | | | 1-5 vs 15-20 tones | | | | |
| | | | Control | 74,58 | 44.30 to 104.9 | *** | 0,0004 |
| | | | NSC 23766 | 91,65 | 69.01 to 114.3 | **** | <0.0001 |
| | | | SCH 23390 | 58,14 | 39.21 to 77.07 | *** | 0,0001 |
| | | | SULPIRIDE | 66,87 | 27.33 to 106.4 | ** | 0,0051 |
| S2E | Days for Crossing the Threshold | The drugs do not have effect itself over the extinction criterion | 25% Percentile | 4 | 3,75 | 3 | 3,75 |
| | | | Median | 4 | 5,5 | 4 | 4,5 |
| | | | 75% Percentile | 5,5 | 6,25 | 4,25 | 5 |
| | | | Kruskal-Wallis test | | | | |
| | | | P value | 0,2843 | | | |
| | | | Kruskal-Wallis statistic | 3,797 | | | |
| | | | 25% Percentile | 12,53 | 26,27 | 13,63 | 10,87 |
| | | | Median | 16,73 | 36,8 | 20,33 | 20,47 |

| Figure | Panel Title | Conclusion | Supporting Data & Statistics | | | | |
|---|---|---|---|---|---|---|---|
| S2F | Renewal Test | During the renewal, NSC2376 seems to increase freezing behavior only compared to control animals infused with saline | 75% Percentile | 20,93 | 52,27 | 26,93 | 25,43 |
| | | | Kruskal-Wallis test | | | | |
| | | | P value | 0,0125 | | | |
| | | | Kruskal-Wallis statistic | 10,86 | | | |
| | | | Dunn's multiple comparisons test | Mean rank diff. | Summary | Adjusted P Value | |
| | | | Control vs. NSC 23766 | -12,5 | * | 0,0132 | |
| | | | Control vs. SCH 23390 | -2,917 | ns | >0.9999 | |
| | | | Control vs. SULPIRIDE | -2,583 | ns | >0.9999 | |
| | | | NSC 23766 vs. SCH 23390 | 9,583 | ns | 0,1133 | |
| | | | NSC 23766 vs. SULPIRIDE | 9,917 | ns | 0,0907 | |
| | | | SCH 23390 vs. SULPIRIDE | 0,3333 | ns | >0.9999 | |
| | | | Test details | Mean rank 1 | Mean rank 2 | Mean rank diff. | |
| | | | Control vs. NSC 23766 | 8 | 20,5 | -12,5 | |
| | | | Control vs. SCH 23390 | 8 | 10,92 | -2,917 | |
| | | | Control vs. SULPIRIDE | 8 | 10,58 | -2,583 | |
| | | | NSC 23766 vs. SCH 23390 | 20,5 | 10,92 | 9,583 | |
| | | | NSC 23766 vs. SULPIRIDE | 20,5 | 10,58 | 9,917 | |
| | | | SCH 23390 vs. SULPIRIDE | 10,92 | 10,58 | 0,3333 | |
| S2G | Reinstatement Test | No differences were detected during the reinstatement test | 25% Percentile | 22,4 | 25,27 | 30,83 | 23,2 |
| | | | Median | 32,4 | 32,2 | 33,07 | 40,27 |
| | | | 75% Percentile | 40,77 | 69,33 | 47,9 | 53,83 |
| | | | Kruskal-Wallis test | | | | |
| | | | P value | 0,7594 | | | |
| | | | Kruskal-Wallis statistic | 1,173 | | | |
| S3A | Test After Training | No difference in fear expression in response to the CS+ following training | Number of Animals | 6 | 6 | | |
| | | | | **Open-Loop** | **Closed-loop** | | |
| | | | 25% Percentile | 65,67 | 50,27 | | |
| | | | Median | 77,73 | 72,13 | | |
| | | | 75% Percentile | 85,63 | 97,3 | | |
| | | | Mann Whitney test | | | | |
| | | | P value | 0,6991 | | | |
| | | | Sum of ranks in column A,B | 42 , 36 | | | |
| | | | Mann-Whitney U | 15 | | | |
| S3B | Transition to Extinction | All groups are able to decrease the fear response over time | 25% Percentile | 14,73 | 10,05 | | |
| | | | Median | 37,22 | 51,43 | | |
| | | | 75% Percentile | 59,72 | 92,81 | | |
| | | | Mixed-Effect Analysis | P value | P value summary | F (DFn, DFd) | |
| | | | Time Factor (1-5 vs 15-20 tones) | <0.0001 | **** | F (1, 10) = 261.9 | |
| | | | Group Factor | 0,0414 | * | F (1, 10) = 5.473 | |
| | | | Group Factor x Time Factor | 0,0007 | *** | F (1, 10) = 22.90 | |
| | | | Bonferroni's multiple comparisons test | Predicted (LS) mean diff. | 95.00% CI of diff. | Summary | Adjusted P Value |
| | | | 1-5 vs 15-20 tones | | | | |
| | | | Open Loop | 44,99 | 30.29 to 59.69 | **** | <0.0001 |
| | | | Closed Loop | 82,77 | 68.06 to 97.47 | **** | <0.0001 |
| S3C | Days for Crossing the Threshold | Animals exposed to closed-loop stimulation required less extinction sessions to achieve the remission criterion | 25% Percentile | 4,5 | 2 | | |
| | | | Median | 6 | 3 | | |
| | | | 75% Percentile | 6,25 | 3 | | |
| | | | Mann Whitney test | | | | |
| | | | P value | 0,0108 | | | |
| | | | Sum of ranks in column A,B | 55 , 23 | | | |
| | | | Mann-Whitney U | 2 | | | |
| S3D | Renewal Test | Closed-loop neuromodulation induce lower fear expression during renewal | 25% Percentile | 20,93 | 2,633 | | |
| | | | Median | 32,2 | 3,067 | | |
| | | | 75% Percentile | 44,9 | 9,267 | | |
| | | | Mann Whitney test | | | | |
| | | | P value | 0,0022 | | | |
| | | | Sum of ranks in column A,B | 57 , 21 | | | |
| | | | Mann-Whitney U | 0 | | | |
| S3B-E | Reinstatement Test | Closed-loop prevents the fear recovery induced by an immediate foot-shock | 25% Percentile | 27 | 5,233 | | |
| | | | Median | 35,73 | 10,07 | | |
| | | | 75% Percentile | 46,23 | 24,13 | | |
| | | | Mann Whitney test | | | | |
| | | | P value | 0,0152 | | | |
| | | | Sum of ranks in column A,B | 54 , 24 | | | |
| | | | Mann-Whitney U | 3 | | | |
